# A Distributed Circuit for Associating Environmental Context to Motor Choice in Retrosplenial Cortex

**DOI:** 10.1101/2020.12.20.423684

**Authors:** Luis M. Franco, Michael J. Goard

## Abstract

During navigation, animals often use recognition of familiar environmental contexts to guide motor action selection. The retrosplenial cortex (RSC) receives inputs from both visual cortex and subcortical regions required for spatial memory, and projects to motor planning regions. However, it is not known whether RSC is important for associating familiar environmental contexts with specific motor actions. Here, we test this possibility by developing a task in which trajectories are chosen based on the context. We find that mice exhibit differential pre-decision activity in RSC, and that optogenetic suppression of RSC activity impairs task performance. Individual RSC neurons encode a range of task variables, often multiplexed with distinct temporal profiles. However, the responses are spatiotemporally organized, with task variables represented along a posterior-to-anterior gradient along RSC during the behavioral performance, consistent with histological characterization. These results reveal an anatomically-organized retrosplenial cortical circuit for associating environmental contexts to appropriate motor outputs.

## INTRODUCTION

The ability to navigate through familiar locations is a fundamental cognitive capability underlying a wide range of animal behaviors, including foraging, nesting, and escape from predation. A wealth of research has focused on navigational strategies that require a “cognitive map” of the local environment (Burgess et al., 2001; O’Keefe and Dostrovsky, 1971). However, in familiar environments, animals often use route-following strategies that involve recognizing learned environmental contexts and using them to guide motor action. For instance, when traveling between two familiar locations (e.g., home and work), we do not need to rely on a cognitive map, but can instead recognize familiar visual scenes (e.g., particular intersections) and use them to guide our actions. Although this ability seems simple, it requires that neuronal populations integrate diverse streams of sensory cues and learned associations in order to select appropriate motor actions. Despite the importance of this ability for animal behavior, the neural circuitry linking the association of the current environmental context to motor selection is poorly understood.

Subcortical regions such as the hippocampal formation are known to be important for spatial navigation. However, it has become increasingly clear that neocortical regions also play a crucial role in navigation, particularly for remote spatial memories (Miller et al., 2014; Mitchell et al., 2018; Smith et al., 2018; Todd et al., 2019; Vann et al., 2009). In fact, although patients with hippocampal lesions have difficulty learning to navigate novel environments, they retain the ability to navigate through locations that were familiar prior to the lesion (Teng and Squire, 1999). In this regard, the retrosplenial cortex (RSC) has attracted particular attention, due to its anatomical connectivity with visual cortex, parietal cortex, premotor cortex, anterior thalamic nuclei, and the hippocampal formation (Mitchell et al., 2018; Vann et al., 2009). Damage to RSC in humans results in navigation deficits even in familiar locations (Aguirre and D’Esposito, 1999; Mitchell et al., 2018). Similar results have been found in rodents, though the effect appears to vary with the extent of the RSC lesion (Sutherland et al., 1988; Vann and Aggleton, 2004, 2005; Vann et al., 2003; Whishaw et al., 2001). These studies indicate a role for RSC in the storage and recognition of familiar locations during navigation.

Previous studies have shown that subsets of RSC neurons are tuned to allocentric and egocentric spatial variables including position (Alexander and Nitz, 2015; Mao et al., 2017, 2018), head direction (Chen et al., 1994; Clark et al., 2010; Jacob et al., 2017), proximity to landmarks (Alexander and Nitz, 2017; Fischer et al., 2020; Mao et al., 2020; Miller et al., 2019; Powell et al., 2020; Vedder et al., 2017), and proximity to boundaries (Alexander et al., 2020; Laurens et al., 2019; van Wijngaarden et al., 2020). However, RSC neurons are also important for the memory of environmental “contexts”, defined as general locations that are identifiable by invariant sensory features and that are independent of the current position or trajectory (e.g., a specific intersection, regardless of which lane one is in). Specifically, exposure to new environmental contexts increases immediate early gene expression in RSC (Czajkowski et al., 2014; Maviel et al., 2004; Milczarek et al., 2018), and disruption of RSC can impair the recall of specific contexts (Buckley and Mitchell, 2016; Corcoran et al., 2011). Additionally, conditioned freezing responses to a particular environmental context can be artificially evoked via optogenetic stimulation of subsets of RSC neurons that exhibited elevated c-Fos expression during exposure to the context (Cowansage et al., 2014; De Sousa et al., 2019). Taken together, these results suggest a role for RSC in remote memory of environmental contexts independent of egocentric or allocentric spatial variables (Mitchell et al., 2018).

To investigate whether RSC plays a role in associating familiar environmental contexts to motor choice independent of local spatial variables, we developed a virtual T-maze task for head-fixed mice in which the wall pattern determined which turn direction would be rewarded. To dissociate the environmental context from other navigational cues (such as position, orientation, and proximity of boundaries), we decoupled mouse position in the environment from physical locomotion, limiting mice to a binary choice of maze trajectory, which was selected using a rotary joystick. This task design is similar to navigation tasks performed by humans in vehicles or virtual environments. Widefield calcium imaging of the dorsal cortical surface of mice performing the task revealed higher RSC activity prior to decision on correct trials relative to incorrect trials. Photoinhibition of RSC excitatory neurons during the pre-decision period led to impaired performance on the task, indicating that RSC activity plays an important role in the association of environmental contexts to appropriate motor choice. Two-photon imaging of populations of RSC neurons indicated that individual neurons encode sensory context, motor choice, and trial outcome variables, with a subset of neurons jointly encoding multiple variables with distinct temporal dynamics. Finally, we found anatomical gradients in the encoding of particular task variables, with posterior RSC encoding environmental context and choice bias signals, while motor planning, motor execution, and trial outcome were encoded throughout the extent of RSC. This functional organization is consistent with anatomical connectivity from visual cortex to posterior RSC and from anterior RSC to forelimb motor area, respectively. Taken together, these results reveal a circuit spanning posterior-to-anterior RSC for associating environmental contexts to appropriate motor planning signals, even in the absence of physical locomotion.

## RESULTS

### Association of environmental context to trajectory in the absence of locomotion

To investigate how the cortex encodes decisions using environmental contexts in the absence of physical locomotion, we developed a virtual T-maze task in which head-fixed mice use a rotary joystick to select one of two trajectories based on visual cues (**Figure 1A**; **Video S1**; **Figure S1**; see Methods). The contexts varied with respect to pattern (context 1: dashed; context 2: striped), orientation (context 1: 90° from vertical; context 2: 45° from vertical), and color (context 1: yellow; context 2: blue). Each context was associated with a particular rewarded arm (**Figure 1B**; context 1: left; context 2: right). Note that the contexts were symmetrical on both sides of the maze, so mice had to recall associations to abstract visual patterns rather than simply orienting to a stimulus appearing on one side. At the start of each trial, the joystick was locked to the center position, and the mouse was presented with one of the two contexts. The mouse then progressed automatically at a fixed speed through the stem of the virtual T-maze (3 s duration) until stopping at a decision point shortly before the split in the T-maze. At this point, the joystick was unlocked, and mice were given a 3 s window to rotate the joystick left or right to a set angular threshold to indicate their decision. During the decision window, the joystick movements were locked to rotation of the virtual environment in a closed loop. If the mouse made a decision within the window, the joystick was locked and a sound indicating the trial outcome was played. The mouse then proceeded automatically to the end of the selected T-maze arm (3 s duration), which contained a visual cue indicating reward for correct decisions or no cue for incorrect decisions. Finally, there was a fixed 3 s inter-trial interval before mice were teleported back to the beginning of the T-maze for the start of the next trial.

**Figure 1.**
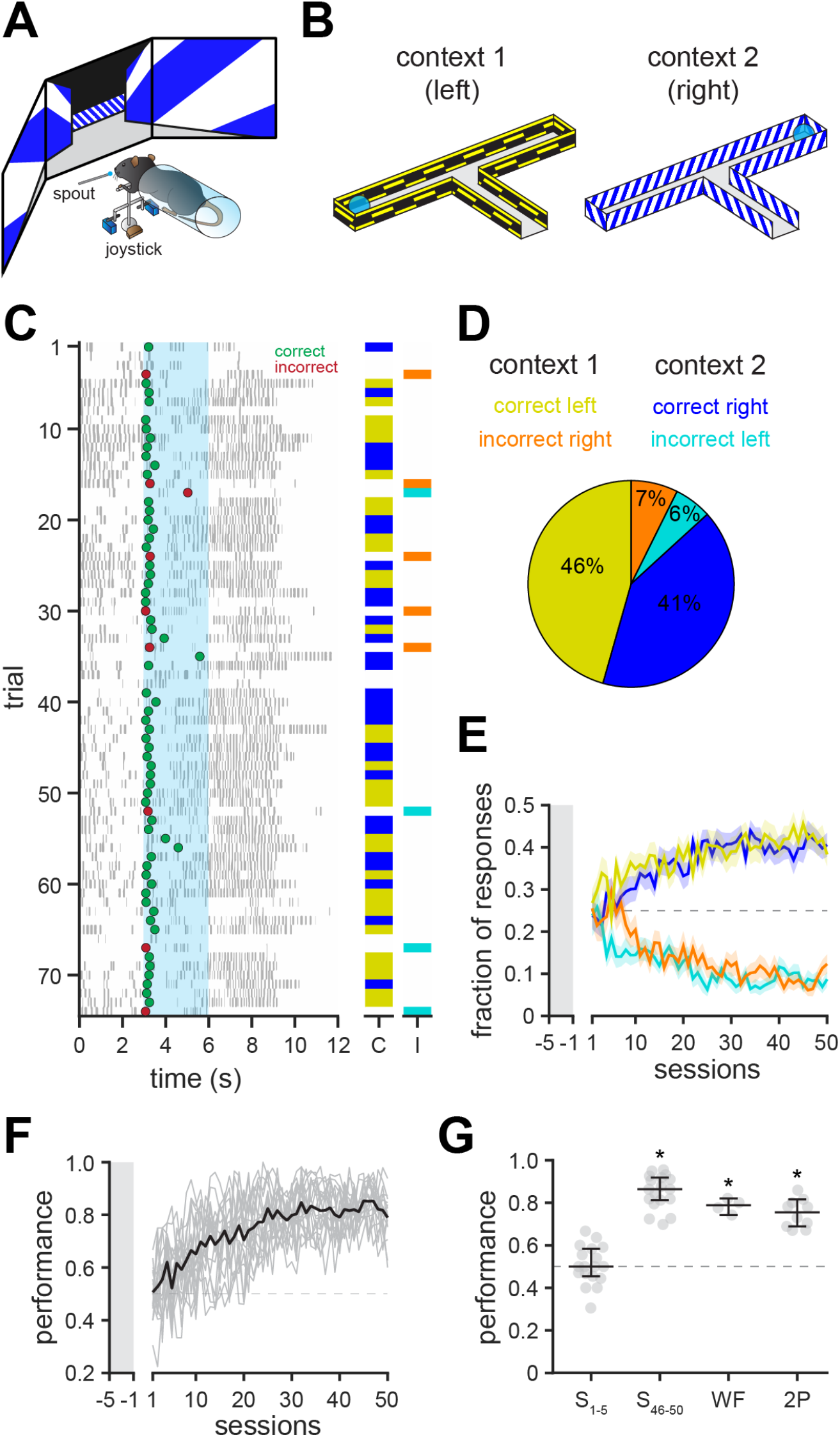
Association of context to trajectory in the absence of physical locomotion. **A**. Schematic of behavioral setup. Mice use a rotary joystick to choose trajectories in a virtual T-maze presented on monitors spanning the horizontal visual field. **B**. Schematic of T-maze task for head-fixed mice in the absence of locomotion. Mice travel at a fixed speed through the stem of the T-maze before stopping shortly before the junction of the maze. At this point, the joystick is unlocked by servomotors and mice must turn the joystick depending on the context. Left turns are rewarded in context 1 (horizontal yellow dashes on black background) whereas right turns are rewarded in context 2 (oblique blue stripes on white background). If a response is registered, the joystick is locked in the center position, and mice proceed to the end of the maze, where a reward (water, 10 μL) is provided if the decision was correct. **C**. Example experimental session showing the behavioral performance of a single mouse. Dots indicate the time at which a decision was registered (green = correct, red = incorrect) or the licking of the water spout (grey ticks); light blue shading indicates the decision window. Columns of the right show correct (C) incorrect (I) responses in context 1 (yellow and orange) and context 2 (blue and cyan), respectively. Trials in which no choice was made within the decision window are left blank. **D**. Summary of the behavioral responses for the session shown in C (yellow = correct left, orange = incorrect right, blue = correct right, cyan = incorrect left). **E**. Fraction of correct and incorrect responses during behavioral training (*n* = 18 mice). Mice are first habituated to the training rig for 5 days (grey shaded area). During this period, the joystick is unilaterally prevented from rotating in the incorrect direction to help mice learn the correct context-reward associations. Following habituation, mice are progressively trained to perform greater numbers of free decisions (10-80% of trials). Only free decision trials are analyzed. Lines and shaded areas show mean ± s.e.m. **F**. Performance during behavioral training (*n* = 18 mice). Performance of individual mice is shown in grey. Average performance is shown in black. **G**. Scatter plots displaying performance of individual mice during the first 5 days of training (S1-5), last 5 days of training (S46-50), widefield imaging sessions (WF) and 2-photon imaging sessions (2P). Individual mice are represented by grey dots. Mouse performance was significantly above chance at the end of training and imaging sessions (S_1-5_ = 50.0, 95% CI = [45.5%, 58.3%]; S_46-50_ = 86.4, 95% CI =[81.3%, 91.9%]; WF = 78.9, 95% CI = [74.2%, 82.0%]; 2P = 75.5, 95% CI = [67.1%, 81.6%]; significant if lower 95% confidence interval > 50%). Black overlay: Median ± 95% CI.

After 46 - 50 training sessions, mice (*n* = 18) increased the number of responses up to an average of 81.1 ± 2.9 decisions per session with a mean performance rate of 82.5 ± 0.8% (**Figure 1C-F**, **Figure S2A, B**). Learning was accompanied by a decrease in response latency (**Figure S2C, D**) and a selective increase in correct decisions (**Figure S2E, F**). We observed no consistent bias in left versus right responses throughout training (**Figure S2G**). Performance was similar between expert training sessions (82.5 ± 0.8%), widefield imaging sessions (78.5 ± 1.5%), and 2-photon imaging sessions (74.3 ± 1.4%; **Figure 1G**). To determine whether mice waited until the decision point before planning their response, we transferred a subset of fully trained mice to a closed-loop version of the task in which forward movement progressed automatically, but angular rotation was controlled in a closed-loop fashion throughout the T-maze. We found that mice performed significantly above chance (72.4 ± 1.4%), and began turning toward the chosen direction well before the decision point (**Figure S2H-K**), indicating that choice planning was initiated prior to the decision window. Note that mice with full control of their trajectory would have exhibited different head direction, proximity to boundaries, and visual input across the trials types, conflating the environmental context with other sensory and spatial variables.

### Mesoscale cortical contributions to accurate context-to-trajectory associations

As a first step toward examining cortical responses during the association task, we imaged the dorsal cortex in transgenic mice with pan-excitatory expression of GCaMP6s (*n* = 20 imaging sessions in 4 mice; **Figure 2A**; see Methods). For these mice, the skull was polished and overlaid with a coverslip to allow imaging of an 8 - 10 mm diameter region, including visual (VC), retrosplenial (RSC), parietal (PPC), somatosensory (SSC), and somatomotor (SMC) cortices (**Figure 2B, C**). During all trial types, activity was highest in VC and RSC prior to the decision (0 - 3 s; **Figure 2D**). Following the joystick movement (3 - 6 s; **Figure 2D**), activity was highest in the SMC and anterior SSC, centered over the caudal forelimb area (CFA), which is necessary for planning and executing forelimb movements (Morandell and Huber, 2017; Tennant et al., 2011). To determine if particular cortical regions exhibited enhanced activity during successful context-to-trajectory associations, we measured the subtracted response (correct trials - incorrect trials), revealing a small but significant increase in activity in RSC, but not other regions during the pre-decision period (**Figure 2E, G; Figure S3A, B**). The higher RSC activity slightly preceded the trial start (−1.5 - 0 s window; **Figure 2E, G**), indicating that the higher activity may be due to higher behavioral engagement on correct trials in addition to differences in encoding of task variables. After the decision point, activity was higher in several regions on correct trials, most noticeably the caudal forelimb area (CFA) (Tennant et al., 2011), spanning anatomically defined SSC and SMC regions (**Figure 2F, G**). Moreover, space-frequency singular value decomposition analysis (Makino et al., 2017) revealed notable differences in the initiation and spatiotemporal progression of neural activity between correct and incorrect trials (**Figure S3C**).

**Figure 2.**
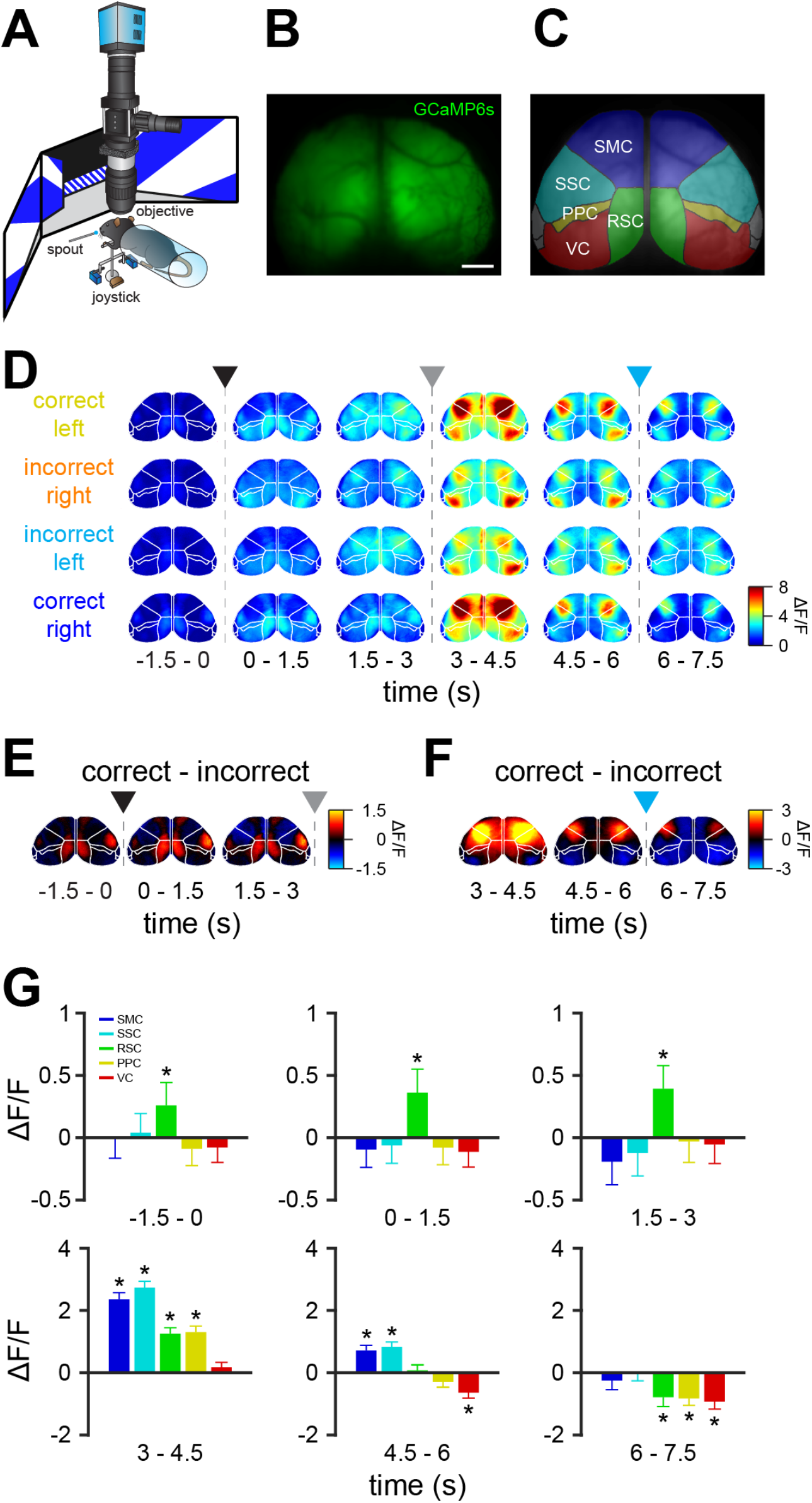
Differential mesoscale activity in RSC during context-trajectory associations. **A**. Schematic of widefield imaging of calcium activity across dorsal cortex during task performance. **B**. The skull of transgenic mice expressing GCaMP6s was polished and overlaid with cyanoacrylate and a coverslip to allow imaging of an 8 - 10 mm diameter region of dorsal cortex. Scale bar = 1 mm. **C**. Regions of dorsal cortex imaged during performance of the task (SMC = somatomotor cortex, SSC = somatosensory cortex, RSC = retrosplenial cortex, PPC = posterior parietal cortex and VC = visual cortex). Region parcellation based on Allen Mouse Brain Common Coordinate Framework (Wang et al., 2020b). **D**. Average registered activity maps obtained in 1.5s epochs during traversal of the T-maze for all trial types (*n* = 20 sessions in 4 mice). Arrowheads and dashed lines indicate trial start (black), decision point (grey) and trial end (cyan). **E-F**. Maps obtained by subtracting correct trials minus incorrect trials. Note that activity is higher in RSC in correct trials prior to the decision point (pooled across both contexts). After decisions, activity is higher in SMC in correct trials. Arrowheads and dashed lines indicate trial start (black), decision point (grey) and trial end (cyan). **G**. Bar plots showing significantly higher RSC activity in correct trials before the decision point (RSC_-1.5-0s_ = 0.26 ± 0.19 ΔF/F, *p* = 0.021; RSC_0-1.5s_ = 0.36 ± 0.19 ΔF/F, *p* = 0.0046; RSC_1.5-3s_ = 0.39 ± 0.19 ΔF/F, *p* = 0.0035). After decisions, SMC and SSC exhibit higher activity (SMC3-4.5s = 2.36 ± 0.21 ΔF/F, *p* = 8.8×10^−8^; SSC_3-4.5s_ = 2.73 ± 0.21 ΔF/F, *p* = 3.6×10^−8^; SMC_4.5-6s_ = 0.71 ± 0.17 ΔF/F, *p* = 3.0×10^−4^; SSC_4.5-6s_ = 0.83 ± 0.15 ΔF/F, *p* = 2.9×10^−5^), whereas RSC and PPC have both higher (RSC_3-4.5s_ = 1.25 ± 0.19 ΔF/F, *p* = 3.3×10^−6^; PPC_3-4.5s_ = 1.30 ± 0.19 ΔF/F, *p* = 2.4 ×10^−6^) and lower (RS_C6-7.5s_ = −0.79 ± 0.30 ΔF/F, *p* = 0.0139; PPC_6-7.5s_ = −0.83 ± 0.22 ΔF/F, *p* = 0.0011) activity in correct trials. VC activity, by contrast, is lower by the end of correct trials (VC_4.5-6s_ = −0.64 ± 0.18 ΔF/F, *p* = 0.0018; VC_6-7.5s_ = −0.92 ± 0.24 ΔF/F, *p* = 0.0014). All values are mean ± s.e.m., compared by Wilcoxon sign-rank test.

To test whether RSC activity is necessary for performance of the association task, we virally expressed the inhibitory opsin ArchT along the anterior-posterior extent of RSC in both hemispheres (**Figure 3A, B**). Recent reports have suggested that direct photoinhibition can reduce off-target effects compared to indirect photoinhibition via stimulation of inhibitory neurons (Babl et al., 2019; Li et al., 2019). We found that ArchT-mediated suppression was power-dependent and resulted in no detectable off-target suppression of cortical activity outside of RSC (**Figure 3C, D**), though we cannot rule out off-target suppression of interconnected subcortical regions such as the anterior thalamic nuclei (Otchy et al., 2015). During randomly interleaved light exposure trials, we found that RSC inhibition resulted in a substantial impairment in task performance (**Figure 3E-G**). This impairment was not seen in mice expressing GCaMP6s alone with the same illumination protocol (**Figure 3E-G**), ensuring the effect is not due to non-specific effects on tissue temperature (Owen et al., 2019). Taken together, these results indicate that RSC plays an important role during context-to-trajectory associations, even when the mouse is not required to physically navigate through space.

**Figure 3.**
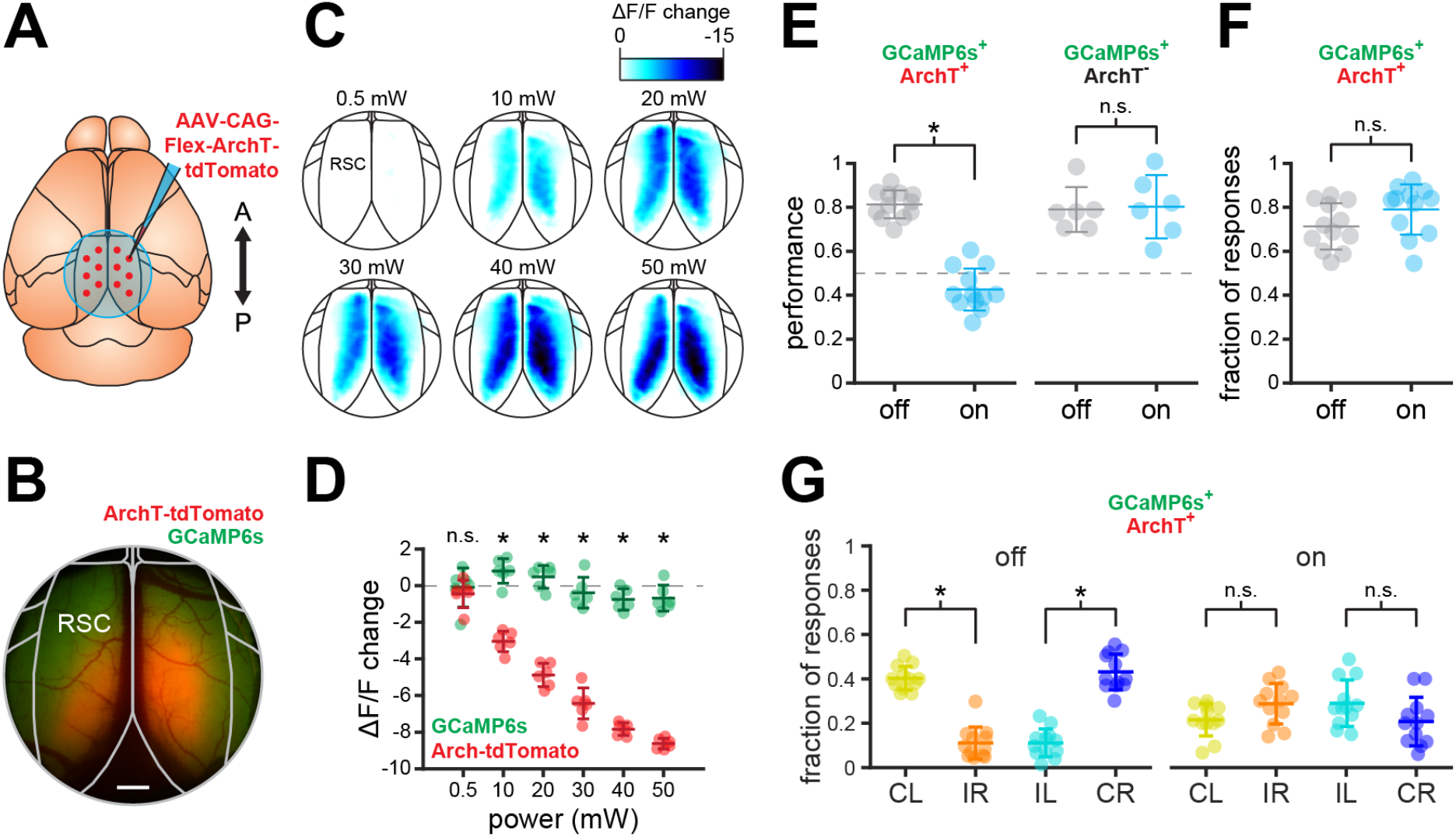
Suppression of RSC activity impairs correct context-trajectory associations. **A**. Transgenic mice expressing GCaMP6s were injected with AAV-CAG-Flex-ArchT-tdTomato (red dots: approximate injection sites) along the rostro-caudal extent of RSC in both hemispheres to drive expression of the inhibitory opsin ArchT. **B**. Example mouse showing expression of ArchT (red) and GCaMP6s (green). Scale bar = 500 μm. **C**. Activity maps showing the suppression of spontaneous activity in RSC immediately following (1 s window) randomly presented light pulses (550 nm, 0.5-50 mW) compared to immediately prior to the pulse (1 s window) in an example ArchT-expressing mouse at different power intensities. **D**. Change in RSC spontaneous activity in the region exhibiting expression of ArchT (red) and in a control region outside of RSC only exhibiting expression of GCaMP6s (green; see Figure 3B). Increasing LED power progressively decreased activity in RSC (n = 12 sessions in 4 mice; ArchT_0.5mW_ = −0.3 ± 0.2 ΔF/F, Control_0.5mw_ = 0.1 ± 0.3 ΔF/F, p = 0.2145; ArchT_10mW_ = −2.8 ± 0.1 ΔF/F, Control_10mW_ = 1.1 ± 0.2 ΔF/F, p = 3.7×10-5; ArchT_20mW_ = −4.7 ± 0.2 ΔF/F, Control_20mW_ = 0.6 ± 0.1 ΔF/F, p = 3.7×10-5; ArchT_30mW_ = −6.3 ± 0.2 ΔF/F, Control_30mW_ = −0.3 ± 0.2 ΔF/F, p = 3.7×10-5; ArchT_40mW_ = −7.7 ± 0.2 ΔF/F, Control_40mW_ = −0.8 ± 0.2 ΔF/F, p = 3.7×10-5; ArchT_50mW_ = −8.4 ± 0.2 ΔF/F, Control_50mW_ = −0.7 ± 0.1 ΔF/F, p = 3.7×10-5; Wilcoxon sign-rank test; mean ± s.e.m.). **E**. Behavioral performance of ArchT^+^ mice (left) was impaired in randomly interleaved trials (25%) by optogenetic inhibition of RSC activity during initial traversal of the stem of the T-maze and the decision window (*n* = 12 sessions in 4 mice; ArchT^+^_off_ = 80.6 ± 6.3%, ArchT^+^_on_ = 42.2 ± 9.5%, *p* = 0.0002; Wilcoxon sign-rank test; mean ± s.d.), but not in ArchT^−^ mice (right; *n* = 6 sessions in 2 mice; ArchT^−^_off_ = 78.3 ± 10.2%, ArchT^−^_on_ = 79.6 ± 14.3%, *p* = 0.7813; Wilcoxon sign-rank test; mean ± s.d.). **F**. No change in the fraction of behavioral responses between trials with and without optogenetic stimulation in ArchT^+^ mice (ArchT^+^_off_ = 70.7 ± 10.5%, ArchT^+^_on_ = 78.3 ± 11.3%, *p* = 0.9966; Wilcoxon signrank test; mean ± s.d.). **G**. Fraction of correct and incorrect responses for both contexts in ArchT^+^ mice during light off trials (ArchT^+^_CL_ = 38.9 ± 5.4%, ArchT^+^_IR_ = 9.7 ± 7.2%, *p* = 0.0002; ArchT^+^_IL_ = 9.7 ± 6.3%, ArchT^+^_CR_ = 41.7 ± 8.0%, *p* = 0.0002; Wilcoxon sign-rank test; mean ± s.d.) and light on trials (ArchT^+^_CL_ = 21.5 ± 7.3%, ArchT^+^_IR_ = 28.8 ± 9.2%, *p* = 0.9478; ArchT^+^_IL_ = 29.0 ± 10.5%, ArchT^+^_CR_ = 20.7 ± 11.0%, *p* = 0.9072; Wilcoxon sign-rank test; mean ± s.d.).

### Single-neuron encoding of context, motor, and outcome variables in RSC

To determine how individual RSC excitatory neurons encode task variables, we used 2-photon microscopy to measure calcium responses from 7,770 Layer 2/3 neurons (29 planes from 10 mice; all planes located 90-155 μm below pia) in bilateral RSC (**Figure 4A-C**; **Figure S4**; see Methods). A majority (5,194 of 7,770 neurons, 66.8%) of individual neurons exhibited significant reliable responses across one of the two correct trial types (see Methods), with a subset (2,818 neurons, 54.3%) responding to a single correct trial type, and the rest (2,376 neurons, 45.7%) responding to both correct trial types, though often with different temporal dynamics (**Figure 4D**; additional example neurons in **Figure S5**). Aligning the population of responses across experiments to trial time (decision window excluded; see Methods), revealed that roughly equal numbers of neurons preferred each correct trial type (**Figure 4E**). As previously observed, RSC neurons exhibited broad but reliable responses (Miller et al., 2019), with peak responses spanning the entire trial duration (**Figure 4E, F**). However, particular epochs of the trial, including the trial start, decision point, and trial end exhibited higher numbers of responsive neurons (**Figure 4G**). Moreover, the response magnitude of the responsive neurons tended to be larger in the later epochs of the trial (**Figure 4H**).

**Figure 4.**
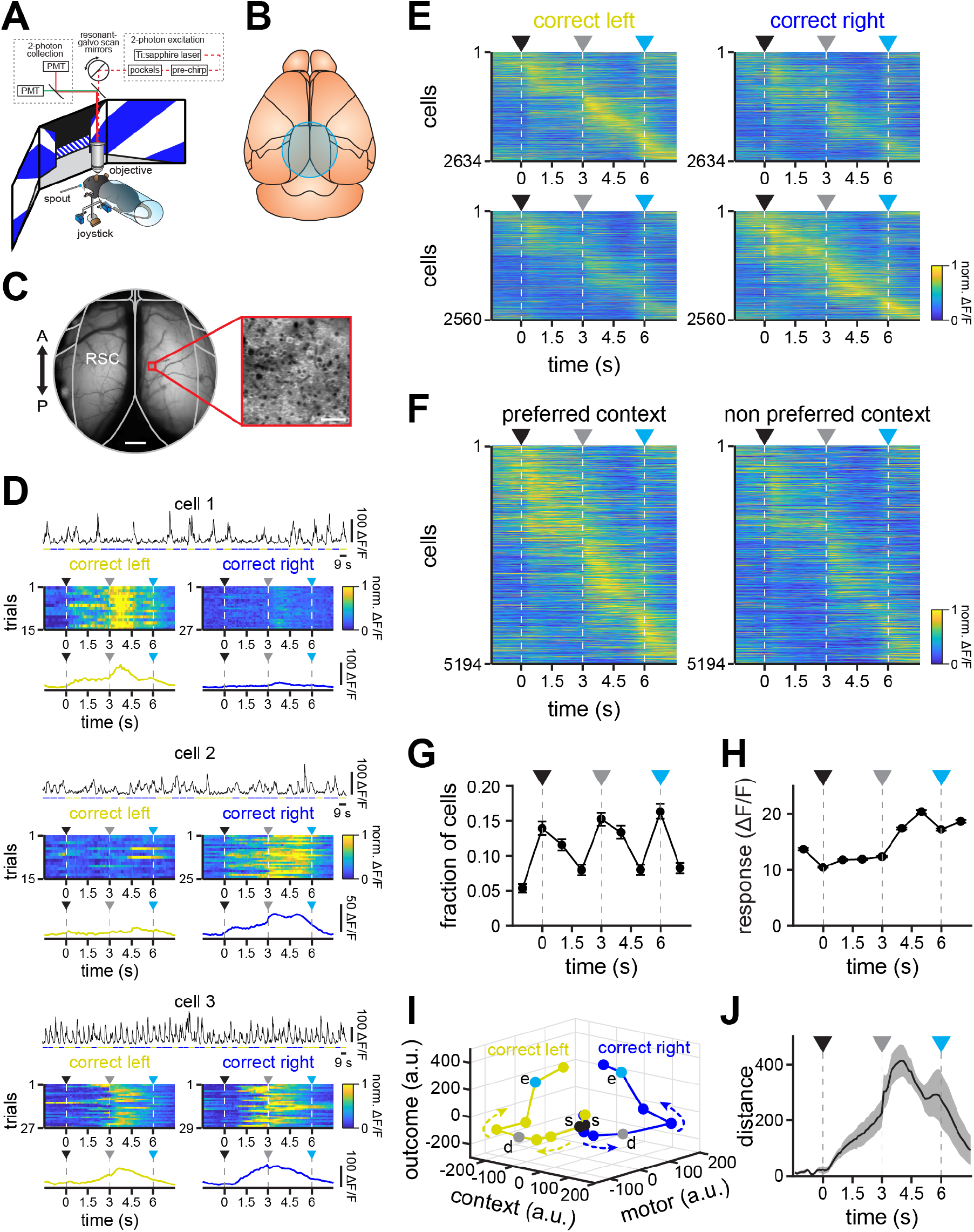
Populations of RSC neurons exhibit selective responses spanning trial duration. **A-B**. Imaging of individual excitatory neurons using 2-photon microscopy (A) through a cranial window over bilateral RSC (B) during performance of the task. **C**. Example of a 4 mm cranial window (left, scale bar = 500 μm) showing an example 225 x 225 μm field (right, scale bar = 50 μm) imaged by 2-photon microscopy. **D**. Example cells exhibiting significant responses for a particular context (cell 1 and cell 2) or for both contexts with different dynamics (cell 3). For each cell: Top, ΔF/F trace across concatenated correct trials for context 1 (yellow) and context 2 (blue). Middle, normalized responses to all correct trials for context 1 (left) and context 2 (right). Bottom, average activity across correct context 1 (left, yellow) and context 2 (right, blue) trials. **E.** Average normalized activity of all neurons selectively responding to correct trials in context 1 (top left, *n* = 2,634 neurons from 10 mice) and in context 2 (bottom right, *n* = 2,560 neurons from 10 mice). Averages only include even trials, with cells sorted by their average latency on odd trials. Activity is also shown for their non-preferred context (top right and bottom left). **F.** Average normalized activity of all significantly responsive neurons for their preferred (left) and nonpreferred (right) context, sorted by peak latency (*n* = 5,194 neurons from 10 mice). Averages only include even trials, with cells sorted by their average latency on odd trials. **G**. Fraction of neurons with peak activity at different times throughout their preferred context (mean ± 97.5 % confidence intervals). **H**. Average responses of all neurons throughout their preferred context (mean ± s.e.m.). **I**. Trajectories of neuronal activity in correct trials for both context obtained by targeted dimensionality reduction (see **Figure S6**). Note that activity is initially similar, however, as trials progress, activity diverges in relation to the particular context and eventual motor decision, followed by a subsequent displacement in the outcome dimension, which is similar for both trajectories (s, trial start; d, decision point; e, trial end). **J**. Distance between trajectories described by neuronal activity in correct trials for both context (mean ± bootstrap-estimated s.e.m.). Arrowheads indicate trial start (black), decision point (grey) and trial end (cyan).

To visualize the population activity in a lower-dimensional space corresponding to task variables, we performed targeted dimensionality reduction (Mante et al., 2013), revealing that different task variables show distinct encoding dynamics in the population throughout the trial (**Figure 4I, J**; **Figure S6**). To further investigate encoding of task variables in individual neurons and populations (Goard et al., 2016; Pho et al., 2018; Pinto and Dan, 2015), we used a linear support vector machine classifier (Raposo et al., 2014; Shahidi et al., 2019) to measure encoding of context (context 1 / context 2), motor output (left / right), and outcome (rewarded / unrewarded) in 100 ms bins spanning the trial (see Methods). A subset of individual neurons encoded single task variables at above-chance levels, though rarely at ceiling levels due to high inter-trial variability (**Figure 5A, B**, cell 1-3; **Figure S7**, cell 5-10). Many individual neurons exhibited multiplexed encoding of multiple task variables with distinct time courses (**Figure 5A, B**, cell 4; **Figure S7**, cell 11-12). The population of RSC neurons was able to decode context, motor output and outcome at high performance levels, though with distinct time courses (**Figure 5C**). Soon after visual cue onset, decoding performance of the context increased and remained high until the offset of visual cues at the end of the trial (**Figure 5C**, top; **Figure 5D, E**). Surprisingly, motor output could be decoded at above-chance levels (62.9 ± 6.0%) prior to visual cue onset, indicating the presence of a motor output bias prior to the start of the trial (**Figure 5C**, middle; **Figure 5D, E**). Motor output encoding performance continued to increase during the approach corridor and peaked shortly after the decision, remaining at high levels throughout the rest of the trial (**Figure 5C**, middle; **Figure 5D, E**). Finally, outcome encoding performance was at chance until after the decision, exhibited a peak during the sound indicating the response outcome, then rose sharply toward the end of the trial (**Figure 5C**, bottom; **Figure 5D, E**). Note that encoding of motor or outcome variables after the decision point may be due to efferent copy feedback resulting from limb movement or licking (Musall et al., 2019; Stringer et al., 2019).

**Figure 5.**
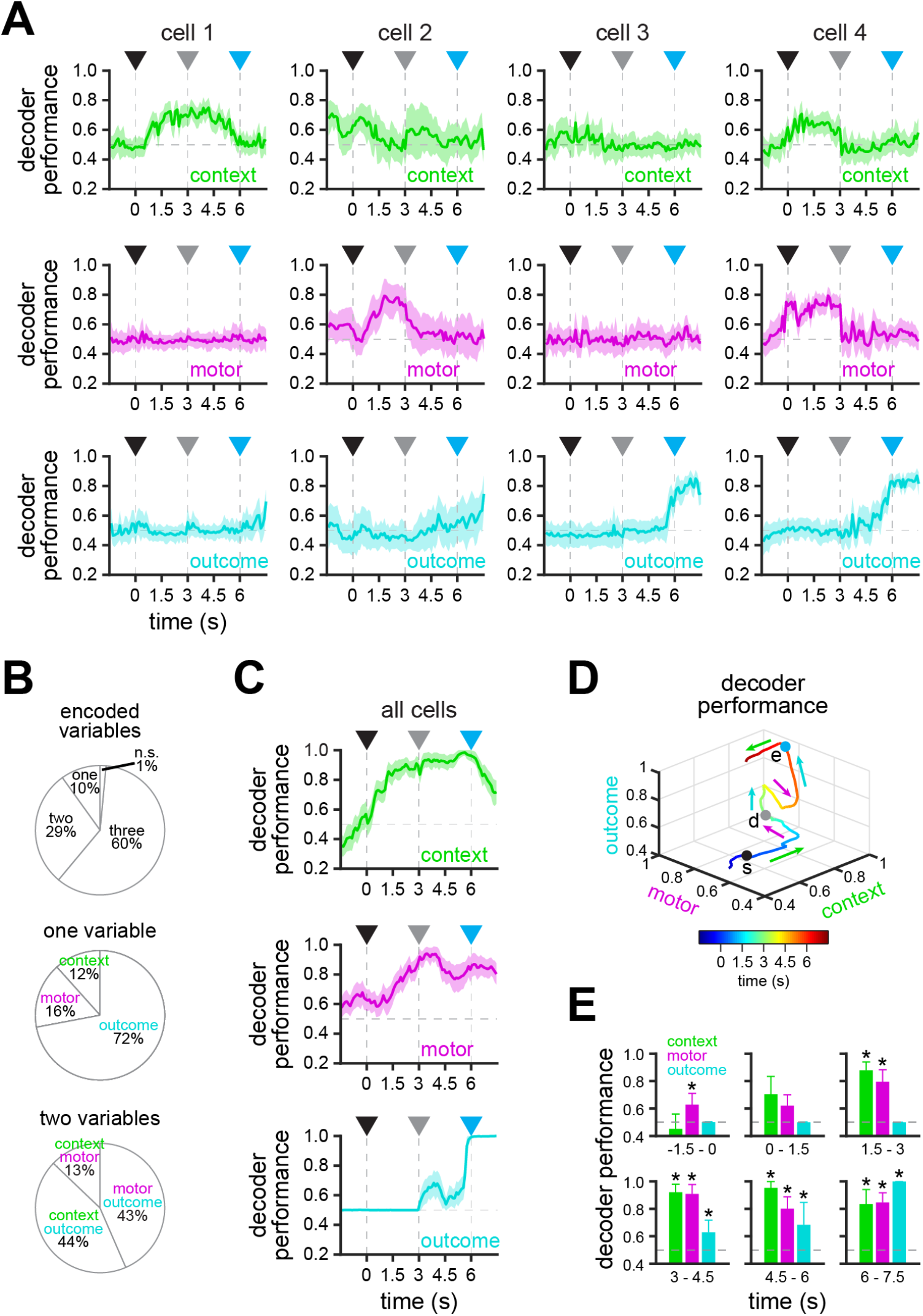
Encoding of task variables by RSC neurons during context-trajectory associations. **A**. Example cells exhibiting significant decoding of a single task variable (cell 1, context; cell 2, motor; cell 3; outcome) or multiple task variables (cell 4) using a support vector machine decoder (chance = 0.5; mean ± bootstrap-estimated s.e.m.). **B**. Top, proportion of cells (*n* = 5,194 from 10 mice) significantly encoding one, two, three or no task variables (see Methods). Middle, cells encoding a single task variable. Bottom, cells encoding a combination of two task variables. **C**. Decoding of context (top), motor (middle) or outcome (bottom) by the entire population of recorded cells (n = 5,194 from 10 mice) combined across sessions using a support vector machine classifier (chance = 0.5; mean ± bootstrap-estimated s.e.m.). **D**. Average trajectory described by population encoding of context, motor and outcome mapped to each axis (s, trial start; d, decision point; e, trial end). Note that encoding of context precedes encoding of the eventual motor action, followed by encoding of outcome. **E**. Bar plots showing significant encoding of the different task variables in different epochs (*n* = 5,194 neurons from 10 mice; lower 5% confidence interval > 0.5; mean ± bootstrap-estimated s.e.m.). Arrowheads indicate trial start (black), decision point (grey) and trial end (cyan).

**Figure 6.**
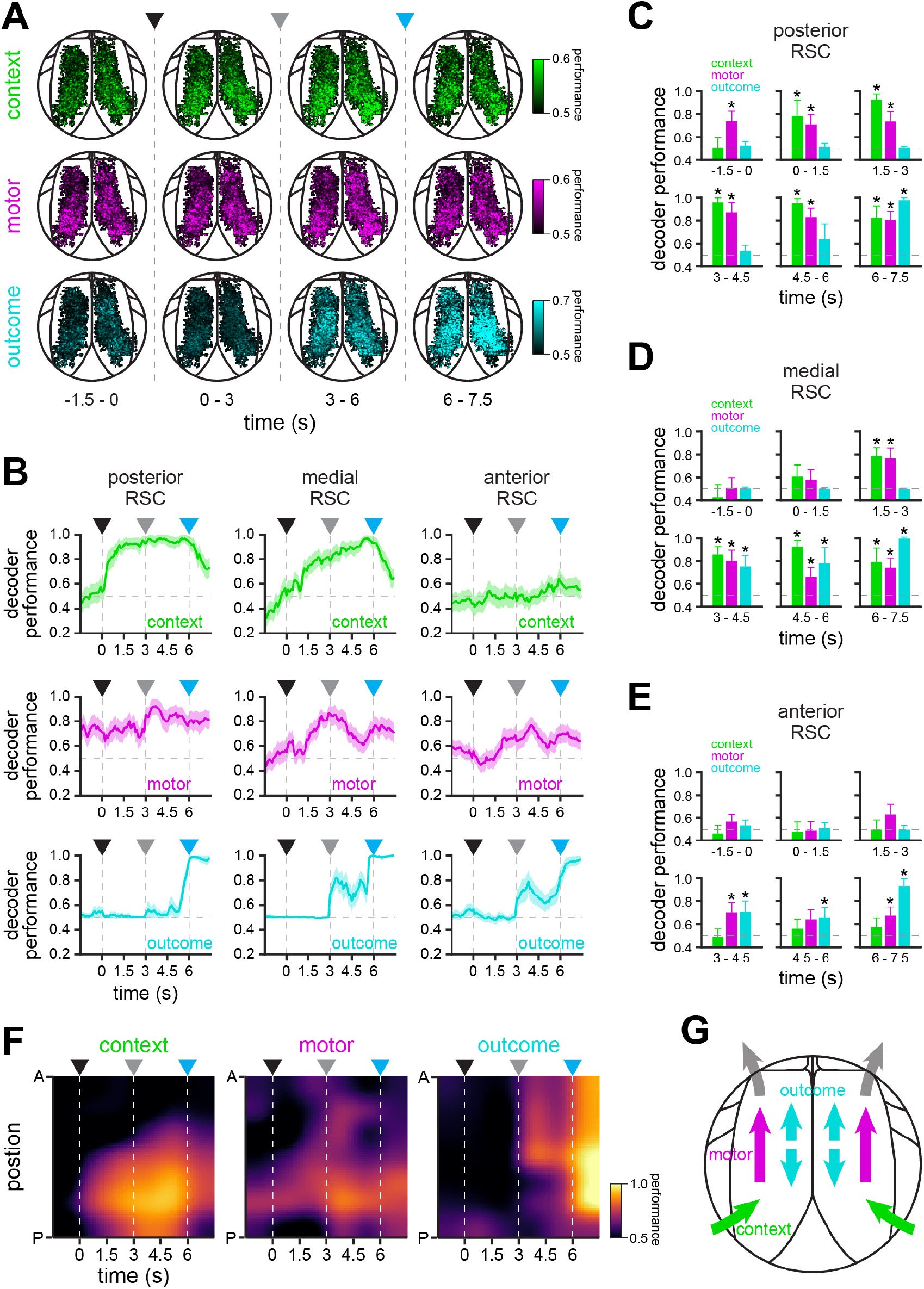
Spatiotemporal encoding of task variables in RSC. **A**. Maps displaying the contribution of individual RSC neurons to the encoding of context, motor and outcome variables across different epochs throughout the trial using a support vector machine classifier (*n* = 5,194 from 10 mice). Some cells are saturated in order to display dynamic range. Note that the population of RSC neurons exhibit different spatial and temporal dynamics. **B**. Encoding of context (top), motor (middle) and outcome (bottom) by different populations of recorded neurons in posterior (*n* = 1,726 neurons), medial (*n* = 2,493 neurons) and anterior (*n* = 975 neurons) RSC (chance = 0.5; mean ± bootstrap-estimated s.e.m.). Note that context is preferentially encoded in posterior RSC, whereas motor and outcome decoding are more distributed across RSC, but with different spatial and temporal dynamics. **C-E**. Bar plots showing significant decoding in posterior (C, n = 1,726 neurons), medial (D, n = 2,493 neurons) and anterior (E, n = 975 neurons) RSC of context, motor and outcome in different task epochs (asterisks: lower 5% confidence interval > 0.5; mean ± bootstrap-estimated s.e.m.). **F.** Spatiotemporal encoding of context, motor and outcome variables by RSC neurons (*n* = 5,194). Context is mainly encoded by neurons in posterior RSC, with higher performance after trial start. By contrast, motor encoding is increased before trial start, revealing pre-trial bias in the eventual motor action. Moreover, motor decoding transitions from posterior to anterior RSC before the decision point and decays shortly before the end of the maze. Outcome is encoded by throughout RSC, with a rapid increase in performance close to the reward location that remains constant until the end of the trial. **G.** Proposed circuit architecture. Environmental context information from visual cortex enters posterior RSC, which then triggers motor planning activity in anterior RSC, including neurons projecting to CFA, the motor region controlling forelimb motor planning and movement. Additionally, outcome information is distributed across all RSC. Arrowheads indicate trial start (black), decision point (grey) and trial end (cyan).

It has recently been shown that trial history is encoded in association cortex (Akrami et al., 2018; Hwang et al., 2017), including RSC (Hattori et al., 2019). Given that motor output encoding was greater than chance prior to trial onset (**Figure 5C**, middle), we suspected that RSC neurons might encode bias from trial history as well as task variables from the current trial. To investigate this, we applied the same approach toward decoding the context, motor output, and trial outcome in the previous trial based on the responses of the current trial. We found that previous-trial context could not be decoded reliably during any epoch of the current trial (**Figure S8**). However, the previous-trial motor output could be decoded with high performance in the pre-trial epoch (−1.5 – 0 s), and with low performance during the pre-decision epoch (0 – 3 s), but could not be decoded during the post-decision epoch (3 – 6 s; **Figure S8**). Finally, previous-trial outcome could be decoded at high performance in the pre-trial and early pre-decision epochs, but not later in the trial (**Figure S8**). This suggests that recent motor output and trial outcome can be decoded from RSC activity (**Figure 5C**, middle; **Figure S8**). However, note that trial history and motor bias on the upcoming trial are related, and cannot be clearly separated with this analysis.

### Spatiotemporal distribution of task variable encoding in RSC

Dysgranular RSC is a large bilateral region in mice, extending several millimeters along the rostrocaudal axis (Mitchell et al., 2018). To determine whether there are hemispheric or anatomical asymmetries in the encoding of task variables, we visualized the encoding performance of all task-responsive neurons organized by spatial position aligned to a reference atlas (**Figure 6A, Video S2**, see Methods). Although encoding was similar across hemispheres (**Figure S9**), task variable encoding exhibited pronounced differences across posterior-to-anterior spatial positions (**Figure 6B-F; Figure S10**). Encoding performance of the context was maximal in the posterior RSC population, weaker in the medial RSC, and virtually absent in the anterior RSC throughout the trial (**Figure 6B**, top row; **Figure 6C-F**). On the other hand, motor encoding performance during the pre-trial period (pre-trial motor bias) was high in the posterior RSC, but as the trial progressed toward the decision point, higher performance was seen in the medial and anterior RSC, indicating a posterior to anterior flow of motor output encoding (**Figure 6B**, middle row; **Figure 6C-F**). Finally, high encoding performance of outcome immediately after the decision point was observed predominantly in the medial and anterior RSC, before becoming widespread shortly prior to the end of the trial (**Figure 6B**, bottom row; **Figure 6C-F**). Results were statistically similar when RSC was divided into different numbers of regions (3-10 regions; data not shown), and when cell number was homogenized across regions by sampling with replacement (**Figure S10**). Together, these results suggest anisotropy along the anterior-posterior axis in the encoding of task variables (**Figure 6G**).

To determine whether this difference in encoding might be due to anatomical differences in cortical inputs and outputs, we performed anterograde and retrograde viral tracer experiments. First, we injected AAV expressing green fluorescent protein in excitatory neurons of the visual cortex (area PM and AM, which project to RSC; Wang et al., 2012) for anterograde labeling of visual cortical axon terminals in RSC (**Figure 7A**). In the same mice, we injected retrograde-transported AAV (Tervo et al., 2016) expressing red fluorescent protein into the caudal forelimb area (CFA) to label RSC neurons projecting to CFA (Morandell and Huber, 2017; Tennant et al., 2011), the primary cortical region active during planning and executing forelimb movements in our task (**Figure 7A**). Both in vivo imaging and histology confirmed that visual cortical inputs predominantly targeted posterior RSC regions, consistent with previous findings (**Figure 7B-D**; Oh et al., 2014; Powell et al., 2020). Conversely, RSC neurons projecting to CFA were located almost exclusively in anterior RSC (**Figure 7B-D**). Together with the posterior-to-anterior asymmetry in spatiotemporal encoding of task variables (**Figure 6F**), these results suggest a simple circuit architecture: visual cues indicating the environmental context originating from visual cortex enter posterior RSC, polysynaptically triggering motor planning activity in anterior RSC, including in neurons projecting to CFA, the motor region controlling forelimb motor planning and movement (schematized in **Figure 6G**).

**Figure 7.**
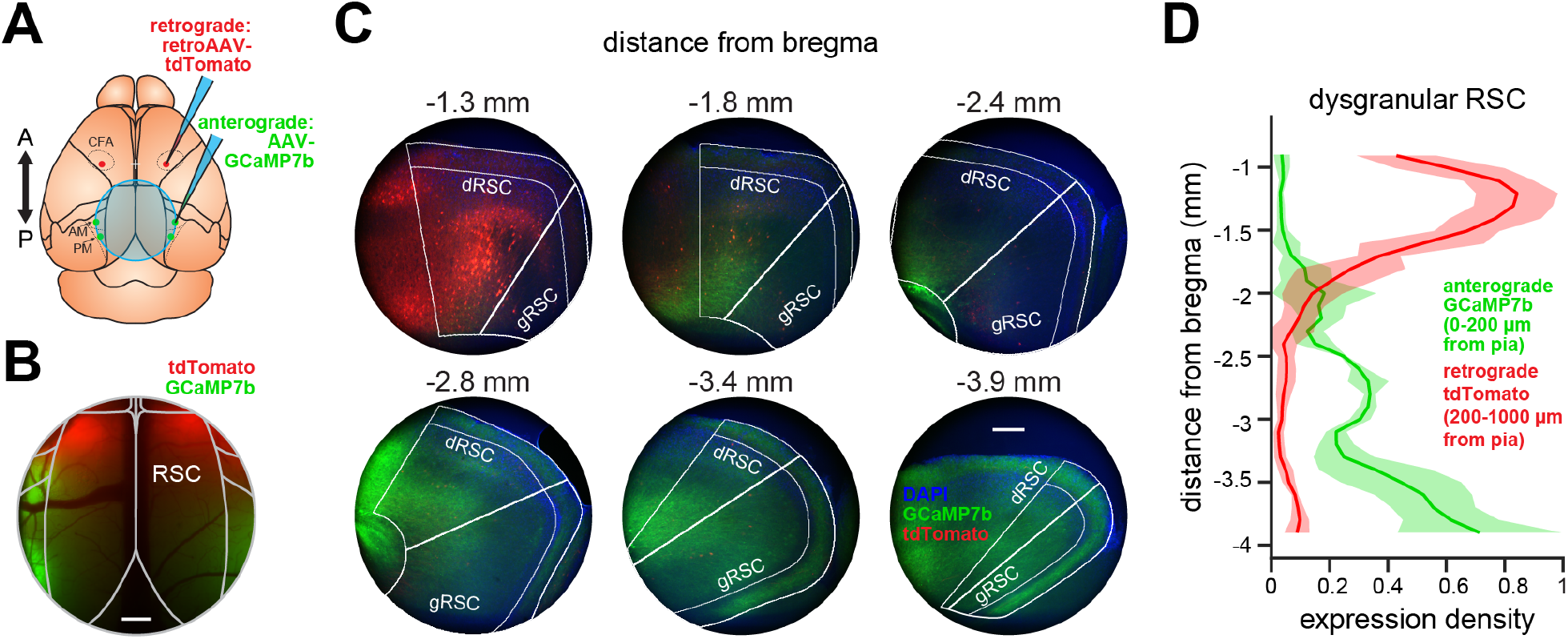
Anatomical gradient of visual inputs and motor outputs in RSC. **A**. Schematic of viral tracing experiments. An anterograde tracer (AAV.Syn.flex.GCaMP7b) was injected bilaterally into regions PM and AM of visual cortex in Emx1-Cre mice to achieve expression in excitatory projections neurons from visual cortex to RSC. A retrograde tracer (AAVrg.CAG.tdTomato) was injected bilaterally into CFA. Finally, a cranial window was implanted over RSC for visualization under the widefield microscope. Cross indicates bregma. **B**. Widefield image of RSC taken 21 days after virus injection. Scale bar = 500 μm. **C**. Example coronal sections taken from anterior (−1.3 mm posterior to bregma) to posterior (−3.9 mm posterior to bregma) of RSC. Axonal expression of anterograde viral tracer (green) is present primarily in posterior RSC while retrogradely labeled cell bodies (red) are almost exclusively located in anterior RSC. Nuclear stain (DAPI, blue) was used for structural visualization. Scale bar = 200 μm). **D**. Normalized expression density (mean ± s.d.) for the anterograde tracer (visual cortical axons, green) in superficial layers (0 - 200 μm from pial surface) and the retrograde tracer (CFA-projecting cell bodies, red) in deep layers (200 - 1000 μm from pial surface) in dRSC. Distributions of expression density are significantly different (*n* = 4 hemispheres from 2 mice; *p* = 0.0046, Kolmogorov–Smirnov test).

## DISCUSSION

Existing theories of RSC function during spatial navigation have posited that RSC is important for three interrelated functions: (1) direct encoding of egocentric and allocentric spatial variables (Miller et al., 2014; Mitchell et al., 2018; Vann et al., 2009), (2) reference frame transformations between allocentric and egocentric representations (Bicanski and Burgess, 2018; Burgess et al., 2001; Mitchell et al., 2018; Vann et al., 2009), and (3) encoding of the general environmental context independent of position (Miller et al., 2014; Mitchell et al., 2018; Smith et al., 2018; Todd et al., 2019). Although a number of recent studies have examined the encoding of spatial variables in RSC (Alexander and Nitz, 2015, 2017; Alexander et al., 2020; Fischer et al., 2020; Jacob et al., 2017; Laurens et al., 2019; Mao et al., 2017, 2018, 2020; Miller et al., 2019; Powell et al., 2020; Vedder et al., 2017; Voigts and Harnett, 2020; van Wijngaarden et al., 2020), much less is known about the role of RSC in encoding general context cues. Coding of environmental context is likely important both for modulating behavioral state, as well as for guiding appropriate reference transformations during decisions. Here, we used a locomotion-independent virtual navigation task to distinguish coding of the environmental context from local spatial variables, such as head direction or proximity to boundaries, which are correlated in freely moving or closed-loop behaviors. This approach revealed the distributed coding of environmental context, motor planning, and outcome in RSC during the association task.

Previous studies have indicated that RSC neurons exhibit weak responses to isolated visual stimuli, such as gratings or objects, even in the context of a behavioral task (Harel et al., 2013; Steinmetz et al., 2019), though visual responses become more robust when they are useful for a spatial task (Fischer et al., 2020). One reason for this may be that visual input to posterior RSC originates primarily from dorsal stream visual areas (Wang et al., 2012), which are specialized for the processing of coherent visual motion (Saleem, 2020; Sit and Goard, 2020), and that RSC responses are particularly responsive to nasal-to-temporal movement (Powell et al., 2020). The RSC may thus be preferentially responsive to whole-field optic flow, textures, and other invariant spatial cues experienced during movement through space, as opposed to playing a general role in visual behaviors. In our study, we used abstract patterns for visual cues instead of unidirectional cues more typically used in virtual T-maze tasks. We reason that while RSC plays an important role for associating abstract features and contexts with appropriate motor responses, detection and orienting responses to unidirectional cues rely more on subcortical structures such as the basal ganglia and superior colliculus (Erlich et al., 2015; Huda et al., 2019; Wang et al., 2020a; Yartsev et al., 2018; Znamenskiy and Zador, 2013).

Although our study focused on RSC, evidence suggests a range of other interconnected regions that also play a role in context-based navigational decisions. Although we see motor planning signals in RSC prior to movement execution (**Figure 5**), similar signals have also been observed in posterior parietal cortex (Driscoll et al., 2017; Pinto et al., 2019), secondary motor cortex (Erlich et al., 2015; Goard et al., 2016; Olson et al., 2020; Pinto et al., 2019), and hippocampus (Gulli et al., 2020). In a recent study investigating the necessity of distributed cortical regions during virtual navigation, only visual cortex was required for an orienting task (turning toward a cue), but a distributed network of visual, parietal, retrosplenial, and motor cortices were required for tasks requiring accumulation or short-term memory of visual cues (Pinto et al., 2019). The coordination of RSC activity with hippocampus, parietal cortex, and motor cortices during navigation requires further study. Pathway specific optogenetic suppression would be one powerful way to approach these questions. It will also be important to further dissect the intra-regional RSC circuit. In particular, what information is carried by posterior-to-anterior RSC projections, and whether silencing of these projections is sufficient to impair the association of context to motor action.

One important finding from our study is the distinct spatiotemporal dynamics of activity representing environmental context and motor planning in RSC. In particular, while posterior RSC encodes context and motor bias resulting from trial history, encoding of motor planning signals moved to medial and anterior RSC prior to the decision (**Figure 6**; **Figure S10**). These results are consistent with increased posterior to anterior flow of widefield calcium responses on correct trials in comparison to incorrect trials (**Figure S3C**), although some motor encoding can be found throughout the RSC. They are also consistent with anatomical data showing that inputs from visual cortex target posterior RSC (Powell et al., 2020; **Figure 7B-D**), while outputs to motor cortex originate from anterior RSC (Yamawaki et al., 2016; **Figure 7B-D**). The anatomical differentiation was even more distinct, indicating that the neurons in posterior RSC encoding the visual context do not directly project to CFA, but must influence motor planning through a polysynaptic pathway. Past work has shown that lesions to posterior RSC preferentially perturb visually-guided spatial navigation, while lesions to anterior RSC disrupts internally guided navigation (Vann and Aggleton, 2005; Vann et al., 2003), further suggesting a functional asymmetry across the posterior-anterior axis. In concordance with this finding, human studies have noted a posterior-to-anterior gradient in RSC, with higher activity in posterior RSC during scene perception, as well as higher activity in anterior RSC during episodic recall and self-referential processing (Chrastil, 2018; Silson et al., 2019). Given the necessity of RSC for the association of environmental context to motor choice, as well as the anatomical differentiation we observed, our results suggest that context is converted into motor commands along the posterior-to-anterior axis of RSC, though we cannot rule out the possibility of cortico-cortico or cortico-striatal projections from posterior RSC influencing motor behavior indirectly. Future studies testing the importance of specific RSC inputs and outputs, as well as projections from posterior to anterior RSC, will be necessary to further dissect this circuit.

## METHODS

### Animals

To achieve widespread calcium indicator expression across dorsal cortex, we bred Emx1-Cre (Jax Stock #005628) x ROSA-LNL-tTA (Jax Stock #011008) x TITL-GCaMP6s (Jax Stock #024104) triple transgenic mice (Madisen et al., 2015) to express GCaMP6s in cortical excitatory neurons. For widefield (*n* = 4), photoinhibition (*n* = 6), 2-photon imaging (*n* = 10), and anatomical tracer (*n* = 2) experiments, 12-16 week old mice of both sexes were implanted with a head plate and cranial window. Water restriction started 7 days after recovery from surgical procedures, and behavioral training started after 7 days of water restriction (14 days after surgery). The animals were housed on a 12 h light/dark cycle in cages of up to 5 animals before the implants, and individually after the implants. All animal procedures were approved by the Institutional Animal Care and Use Committee at University of California, Santa Barbara.

### Surgical Procedures

All surgeries were conducted under isoflurane anesthesia (3.5% induction, 1.5 - 2.5% maintenance). Prior to incision, the scalp was infiltrated with lidocaine (5 mg kg-1, subcutaneous) for analgesia and meloxicam (1 mg kg-1, subcutaneous) was administered pre-operatively to reduce inflammation. Once anesthetized, the scalp overlying the dorsal skull was sanitized and removed, and the periosteum was removed with a scalpel. For widefield imaging experiments, the skull was polished with conical rubber polishing bit, coated with cyanoacrylate (Loctite 406) and overlaid with an 8-10 mm round coverglass. For 2-photon and photoinhibition experiments, the skull was abraded with a drill burr to improve adhesion of dental acrylic. Then, a 4 - 5 mm diameter craniotomy was made over the midline (centered at 2.5 – 3.0 mm posterior to bregma), leaving the dura intact. A cranial window was implanted over the craniotomy and sealed first with silicon elastomer (Kwik-Sil, World Precision Instruments) then with dental acrylic (C&B-Metabond, Parkell) mixed with black ink to reduce light transmission. The cranial windows were made of two rounded pieces of coverglass (Warner Instruments) bonded with a UV-cured optical adhesive (Norland, NOA61). The bottom coverglass (4 mm) fit tightly inside the craniotomy while the top coverglass (5 mm) was bonded to the skull using dental acrylic. A custom-designed stainless steel head plate (eMachineShop.com) was then affixed using dental acrylic. After surgery, mice were administered carprofen (5 – 10 mg kg-1, oral) every 24 h for 3 days to reduce inflammation.

### Virtual T-Maze Design

Mice were head-fixed using custom restraint hardware (https://goard.mcdb.ucsb.edu/resources) and placed in polypropylene tubes to limit movement. A custom rotatory joystick, located within reach of mouse forelimbs, was mounted on optical hardware (Thorlabs) and attached to an optical encoder (Digi-key Electronics). Two servomotors (Adafruit Industries) were used to constrict the position of the joystick throughout the trial, allowing movement only during the decision window. The servos could also be opened unilaterally to allow turning responses in only one direction. A spout was placed near to the mouth of the mouse for delivery of water rewards (10-12 μL), which were controlled by a solenoid valve (Parker Hannifin). Licks were detected through a capacitive touch sensor connected to the metallic spout and to a metallic mesh inside the polypropylene tube. A virtual T-maze was displayed across 3 screens arranged in an arc to subtend 180° of the mouse visual field. All electronic components were controlled by custom code written in Matlab (Mathworks) through Arduino Uno (Arduino).

Virtual mazes were built using the ViRMEn package (Aronov and Tank, 2014), with modifications to control progression through the maze. We created two different virtual T-mazes with unique wall patterns, one consisting of horizontal yellow dashes on a black background (context 1) and one with oblique blue stripes on a white background (context 2). The mazes were presented in pseudorandom order across an experimental session. Progression through virtual environment proceeded at a fixed speed throughout the trial, such that both traversal from the maze start to the decision point and traversal from the decision point to the end of the maze lasted 3 s each. During the decision window, two servomotors opened to allow rotation of the joystick with the forepaws. All mice used both forepaws to rotate the joystick. Rotation of the joystick was coupled to rotation of the field of view in the virtual environment. If the field of view rotation exceeded 45° to the left or to the right from the central position, a decision was registered and the joystick was returned to the central position using the servomotors. If a decision was registered, a tone was played indicating the trial outcome (correct: 5 kHz, 70 dB tone for 0.25 s; incorrect: broadband white noise, 80 dB for 0.25 s), the mouse was automatically rotated 90° in the chosen direction, and then progressed to the end of the selected T-maze arm. The end of the rewarded arm contained a reward cue (a blue sphere) that was reached at the same time as water was administered, whereas the unrewarded arm did not contain the reward cue. As the mouse reached the end of the maze, the outcome tone was played again at the same frequency and volume but longer duration (correct: 5 kHz, 70 dB tone for 1 s; incorrect: broadband white noise, 80 dB for 2 s). If no decision was detected within 3 s, the joystick was returned to the center position using the servos, and the trial was aborted and not included in later analyses. At the end of each trial, there was a 3 s interval with a uniform blue (correct trial) or black (incorrect trial) screen before teleporting back to the start of the T-maze for the next trial. To avoid repeated presentations of the same maze, the probability for consecutively displaying the same context progressively decreased after a particular context was displayed, from 50% to 0% over 10 trials. To prevent bias, if the number responses to the left or to the right exceeded by 50% the number of responses in the non-preferred direction, the context rewarded for the non-preferred direction was consecutively displayed until bias was reduced.

In the closed loop version of the task, mice were allowed to rotate the field of view throughout the approach corridor with a maximum 50° rotation in either direction from the axis of traversal. Forward movement automatically progressed at a fixed speed. Decisions were registered if rotation exceeded 45° to the left or to the right once the mouse reached the decision point, otherwise optical flow stopped for a maximum of 3 s and resumed if a decision was detected.

### Behavioral Training

Before training, mice were gradually water restricted beginning 7 days after surgery. HydroGel (99% H_2_O, ClearH20) was provided in decreasing amounts each day for 7 days (2.0g to 1.2g). During training, mice were supplemented with HydroGel depending on the amount of water obtained in each particular session to keep their body weight at or above 85% of the their initial weight (Goltstein et al., 2018), typically 1.2 mL water (or HydroGel equivalent) per day. In addition, once per week (prior to non-training days), mice received 2.0 g of HydroGel.

Starting 7 days after water restriction, mice spent 5 days of training on a habituation and shaping protocol. During this time, we unilaterally opened the servo to only allow correct decisions (forced choice trials). This stage was important for training mice to move the joystick in both directions without bias, while also instructing them on the correct context-reward associations. Following shaping, the percentage of trials with free choices was progressively increased from 10% to 80%. Only free choices were used for all further analyses. Typically, it took mice 25-40 sessions to achieve plateau levels of performance. One mouse was removed from the study for health reasons prior to completion of training; all other mice were included.

For imaging sessions, in some cases we acquired multiple sessions from the same field of neurons across subsequent days. Since decoding analyses require representation of all trial types (combinations of context and motor choice) and expert mice often make few incorrect choices, we selected imaging sessions *a priori* that contained the highest minimum number of each trial type, with a minimum of 3 trials of each trial type (correct trials, range = 8 - 56 trials, mean ± s.d. = 23.2 ± 9.7 trials; incorrect, range = 3 - 25 trials, mean ± s.d. = 7.8 ± 4.7 trials). Imaging sessions that failed this criterion for any of the four trial types were not included in further analyses. To ensure that our trial threshold did not affect our results, we repeated the analyses with minimum trial thresholds of 4-7 trials. Although the neuron number decreased with more stringent thresholds, the spatiotemporal encoding profiles across task variables were similar (data not shown).

### Widefield Imaging

For widefield imaging experiments (Figure 2), GCaMP6s fluorescence was imaged through the skull (see Surgical Procedures) using a custom widefield epifluorescence microscope (Sit and Goard, 2020) (https://goard.mcdb.ucsb.edu/resources). In brief, broad spectrum (400-700 nm) LED illumination (Thorlabs, MNWHL4) was band-passed at 469 nm (Thorlabs, MF469-35), and reflected through a dichroic (Thorlabs, MD498) to the microscope objective (Olympus, MVPLAPO 2XC). Green fluorescence from the cortex passed through the dichroic and a bandpass filter (Thorlabs, MF525-39) to a scientific CMOS (PCO-Tech, pco.edge 4.2). 400 x 400 pixel frames were acquired at 10 Hz with a field of view of 8.0 x 8.0 mm, leading to a pixel size of 0.02 mm pixel^−1^. A custom light blocker affixed to the head plate was used to prevent light from the visual stimulus monitor from entering the imaging path.

### Widefield Post-processing

Images were acquired with camera control software (pco.edge) and saved into multi-page TIF files. All subsequent image processing was performed in MATLAB (Mathworks). The ΔF/F for each individual pixel of each image frame was calculated as:

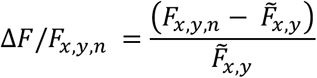

where *F_x,y,n_* is the fluorescence of pixel (*x,y*) at frame *n*, and 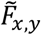 is defined as the median raw fluorescence value across the entire time series for pixel (*x,y*). Subsequent analyses were performed on whole-frame ΔF/F matrices.

In order to define the borders of the different cortical regions for each mouse, we aligned the average projection of the dorsal cortex of each mouse to a template obtained from the Allen Mouse Brain Common Coordinate Framework (Wang et al., 2020b). Briefly, we used custom software written in Matlab (Mathworks) to scale and rotate the template into alignment using the superior sagittal sinus, the transverse sinus, and the borders of the dorsal cortex as reference points.

Space-frequency singular value decomposition was performed using Chronux 2.11 (http://chronux.org). In brief, average ΔF/F time series per mouse corresponding to each pixel were concatenated for correct and incorrect trials, respectively. Slepian sequences were then obtained using 10 tapers (time-bandwidth product = 10), and space-frequency singular value decomposition was calculated for projected pixel time series into frequencies between 0 - 5 Hz. The resulting decomposed time series represent orthogonal sets of spatial modes. The first mode was used as it explained the highest variability in our data (correct trials = 48.02%, incorrect trials = 40.89%). Gradients in the phase of decomposed time series indicate the direction of travelling waves in dorsal cortex during traversal of the virtual T-maze (Makino et al., 2017).

### Photoinhibition Experiments

Prior to cranial window implantation (see Surgical Procedures), a series of 6 - 10 microinjections (50 - 100 nL each, 200-300 μm depth) of AAV5.Flex.ArchT.tdTomato (Addgene, #28305-AAV5) were delivered along the posterior to anterior axis of RSC (2.0 to 4.0 mm posterior, 0.5 mm lateral to bregma) in each hemisphere in triple transgenic GCaMP6s mice. Use of the Emx1-Cre line restricted viral expression of ArchT to excitatory RSC neurons. To characterize cortical inhibition, we performed widefield imaging during randomized periodic flashes (5-10 s) of 0.5 - 50 mW (0.04 – 4 mW m^−2^) light, delivered to the cranial window with a fiber-coupled LED (550 nm, Prizmatix). The widefield ΔF/F immediately after (1 s) light offset was compared to widefield ΔF/F immediately before (1 s) onset. During optogenetics experiments in behaving mice, the RSC activity was suppressed by illumination with 40 mW (~3.2 mW mm^−2^) 550 nm light from the maze onset until decision threshold was reached (3 - 6 s duration). To control for heat-induced changes to neuronal activity or behavior (Owen et al., 2019), we used the same illumination parameters in a set of control mice expressing pan-excitatory GCaMP6s, but not ArchT (light only controls).

### Two-Photon imaging

For 2-photon imaging experiments, GCaMP6s fluorescence was imaged using a Prairie Investigator 2-photon microscopy system with a resonant galvo scanning module (Bruker). Prior to 2-photon imaging, widefield imaging was used to identify the RSC field by alignment to surface blood vessels.

For fluorescence excitation, we used a Ti:Sapphire laser (Mai-Tai eHP, Newport) with dispersion compensation (Deep See, Newport) tuned to λ = 920 nm. For collection, we used GaAsP photomultiplier tubes (Hamamatsu). To achieve a wide field of view, we used a 16x/0.8 NA microscope objective (Nikon) at an optical zoom of 2x (425 x 425 μm field). Imaging planes at a depth of 90-150 μm were imaged at a frame rate of 10 Hz. Laser power ranged from 40-75 mW at the sample depending on GCaMP6s expression levels. Photobleaching was minimal (<1% min^−1^) for all laser powers used. A custom stainless-steel light blocker (eMachineShop.com) was mounted to the head plate and interlocked with a tube around the objective to prevent light from the visual stimulus monitor from reaching the PMTs.

### 2-Photon Post-processing

Images were acquired using PrairieView acquisition software and converted into TIF files. All subsequent analyses were performed in MATLAB (Mathworks) using custom code (https://goard.mcdb.ucsb.edu/resources). First, images were corrected for X-Y movement by registration to a reference image (the pixel-wise mean of all frames) using 2-dimensional cross correlation.

To identify responsive neural somata, a pixel-wise activity map was calculated using a modified kurtosis measure. Neuron cell bodies were identified using local adaptive threshold and iterative segmentation. Automatically defined ROIs were then manually checked for proper segmentation in a graphical user interface (allowing comparison to raw fluorescence and activity map images). To ensure that the response of individual neurons was not due to local neuropil contamination of somatic signals, a corrected fluorescence measure was estimated according to:

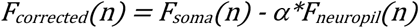

where *F_neuropil_* was defined as the fluorescence in the region within 30 μm from the ROI border (excluding other ROIs) for frame *n* and *α* was chosen from [0 1] to minimize the Pearson’s correlation coefficient between *F_corrected_* and *F_neuropil_*. The ΔF/F for each neuron was then calculated as:

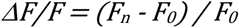

Where *F_n_* is the corrected fluorescence (*F_corrected_*) for frame *n* and *F_0_* defined as the mode of the corrected fluorescence density distribution across the entire time series.

### Analysis of 2-Photon Imaging Data

To determine whether a neuron exhibited consistent task-related activity we calculated the Pearson’s correlation coefficient between odd and even correct trials and compared it to the correlation coefficient of trials in which activity was circularly permuted a random sample number on each trial (time shuffled). If the original correlation coefficient was greater than the 97.5^th^ percentile of the correlation coefficients from the time shuffled activity (1000 iterations), then the neuron was considered to exhibit reliable task-related activity and used for all subsequent analyses.

Targeted dimensionality reduction (Figure 4) was performed as previously described (Goard et al., 2016; Mante et al., 2013). First, neuronal activity was denoised using the first 50 principal components analysis (explained variance = 62.09%). We then used linear regression on the principal components to define de-mixed task-related axes for context, motor choice and outcome. Population trajectories were plotted for correct and incorrect trials in context 1 and context 2 as a function of time throughout the trial. Distances across trajectories were calculated as Euclidean distances in the reduced space for each time bin.

Encoding of task variables (Goard et al., 2016; Raposo et al., 2014) was determined by support vector machine classifiers (Raposo et al., 2014; Shahidi et al., 2019). To achieve this, trials were homogenized by sampling equal number of each trial type with replacement (correct left, incorrect right, incorrect left and correct right) and then split into 2 non-overlapping groups containing 50% of each trial type. This process ensures that all conditions comprise the same number of trials. Then, a support vector machine classifier was trained using 50% of trials and tested on the remaining 50% of trials per time bin. The performance s.e.m. was obtained by taking the standard deviation of bootstrapped samples (100 iterations). Encoding of the different variables was determined by comparing conditions as follows: (1) context: correct left and incorrect right versus incorrect left and correct right, (2) motor: correct left and incorrect left versus incorrect right and correct right, and (3) outcome: correct left and correct right versus incorrect right and incorrect left.

### Anatomical Tracing

For anterograde tracing, we injected 100 μL of AAV1.Syn.Flex.GCaMP7b (Addgene, #104493-AAV1) bilaterally into visual area PM (−3.3 mm posterior, 1.6 mm lateral to bregma) and AM (−2.2 mm posterior, 1.7 mm lateral to bregma) of Slc17a7-Cre mice (Jax Stock #023527) to express a green indicator in excitatory projection axons originating in visual cortex. For retrograde tracing, we made two 100μL injections of AAVrg.CAG.tdTomato (Addgene, #59462-AAVrg) bilaterally into CFA (0.0 mm posterior, 1.6 - 2.0 mm lateral to bregma). We then made a window centered over RSC as described previously. After 2-3 weeks, we performed widefield imaging, and then the animals were perfused with 4% paraformaldehyde. The brain was removed and placed in 4% PFA for 24 h, followed by 24 h in PBS. We then cut 100 μm coronal slices and mounted them on slides with DAPI (Vector Laboratories, #H-1500-10). The slices were imaged on an Olympus BX51 fluorescence microscope and analyzed with Matlab (Mathworks).

### Statistical Information

All data groups were compared using non-parametric tests (Wilcoxon rank-sum, Wilcoxon signed-rank tests or Kolmogorov-Smirnov test). Bootstrap estimates of s.e.m. were calculated as the standard deviation of values evaluated in 100 - 1,000 shuffled iterations, obtained by randomly sampling with replacement from the original values. Differences were considered significant if they were outside the 95^th^ percentile of the bootstrapped distribution.

## DATA AVAILABILITY

All data and code described in this manuscript are available from the lead contact upon reasonable request.

## ACKNOWLEDGEMENTS

We would like to thank Douglas A. Nitz, Abhishek Banerjee, Steffen Kandler, and Dun Mao for comments on manuscript; Tyler Marks for assistance with histology experiments; and Serena Gelfer, Emily Hutchings, Nicole Glick, and Anna Ortiz for assistance with behavioral training. This work was supported by the Harvey Karp Discovery Award (L.M.F.) and UC MEXUS-CONACYT Postdoctoral Fellowship (L.M.F.), NIH R00MH104259 (M.J.G.), NSF 1707287 (M.J.G.), the Hellman Fellows Fund (M.J.G.), the Larry L. Hillblom Foundation (M.J.G.), and the Whitehall Foundation (M.J.G.).

## AUTHOR CONTRIBUTIONS

L.M.F. and M.J.G. designed the experiments; L.M.F. conducted the experiments; L.M.F. and M. J.G. analyzed the data; L.M.F. and M.J.G. wrote the manuscript.

## COMPETING INTERESTS

The authors declare no competing financial interests.

## MATERIALS AND CORRESPONDENCE

Correspondence and material requests should be directed to

## SUPPLEMENTAL INFORMATION

**Figure S1.**
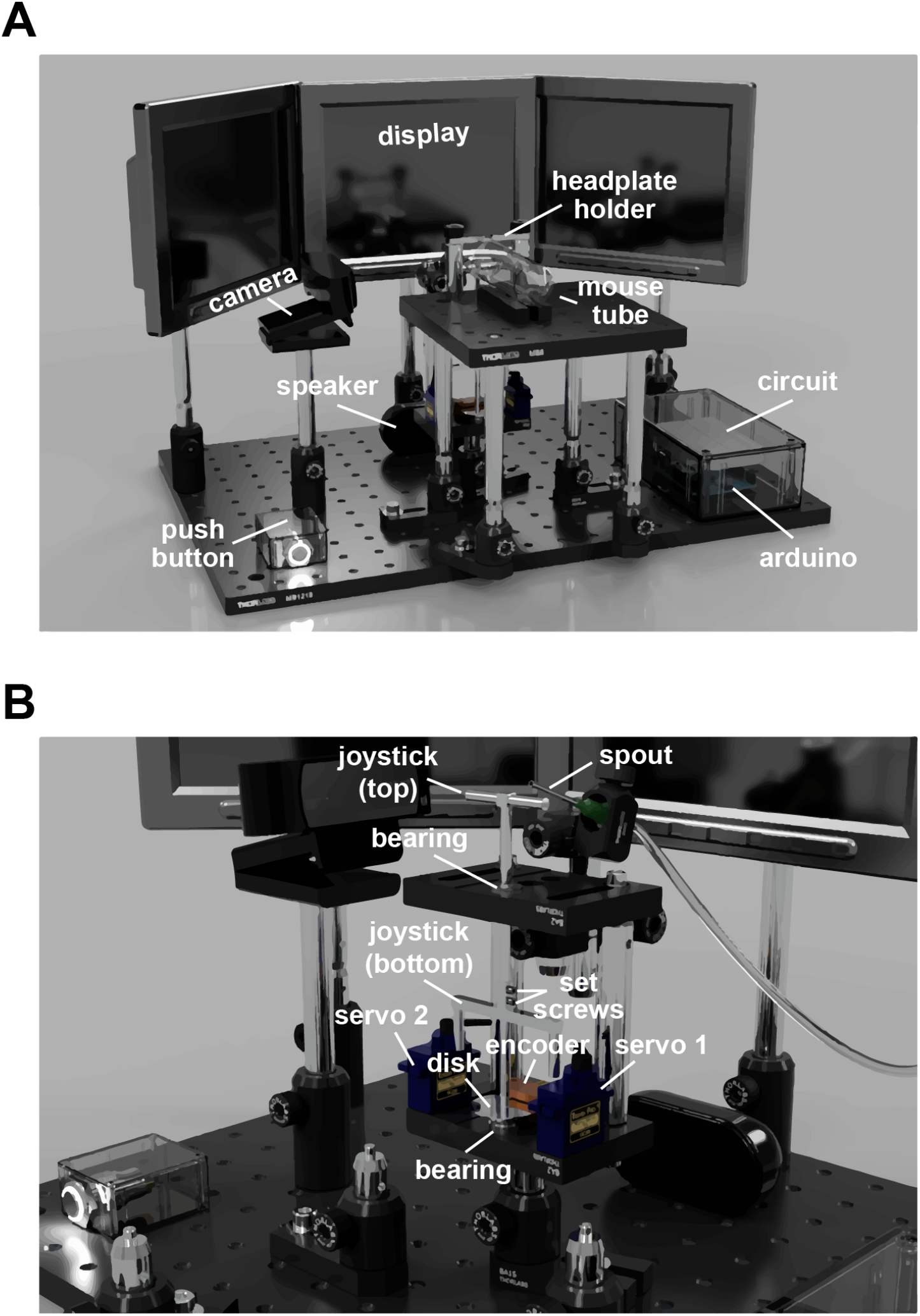
Schematic of behavioral rig hardware. Related to Figure 1. **A**. CAD rendering of the behavior rig used for training mice to perform the context-to-trajectory association task in the absence of locomotion. The location of hardware components is indicated. **B**. Detail of rotary joystick design showing the joystick parts, servomotors and optical encoder.

**Figure S2.**
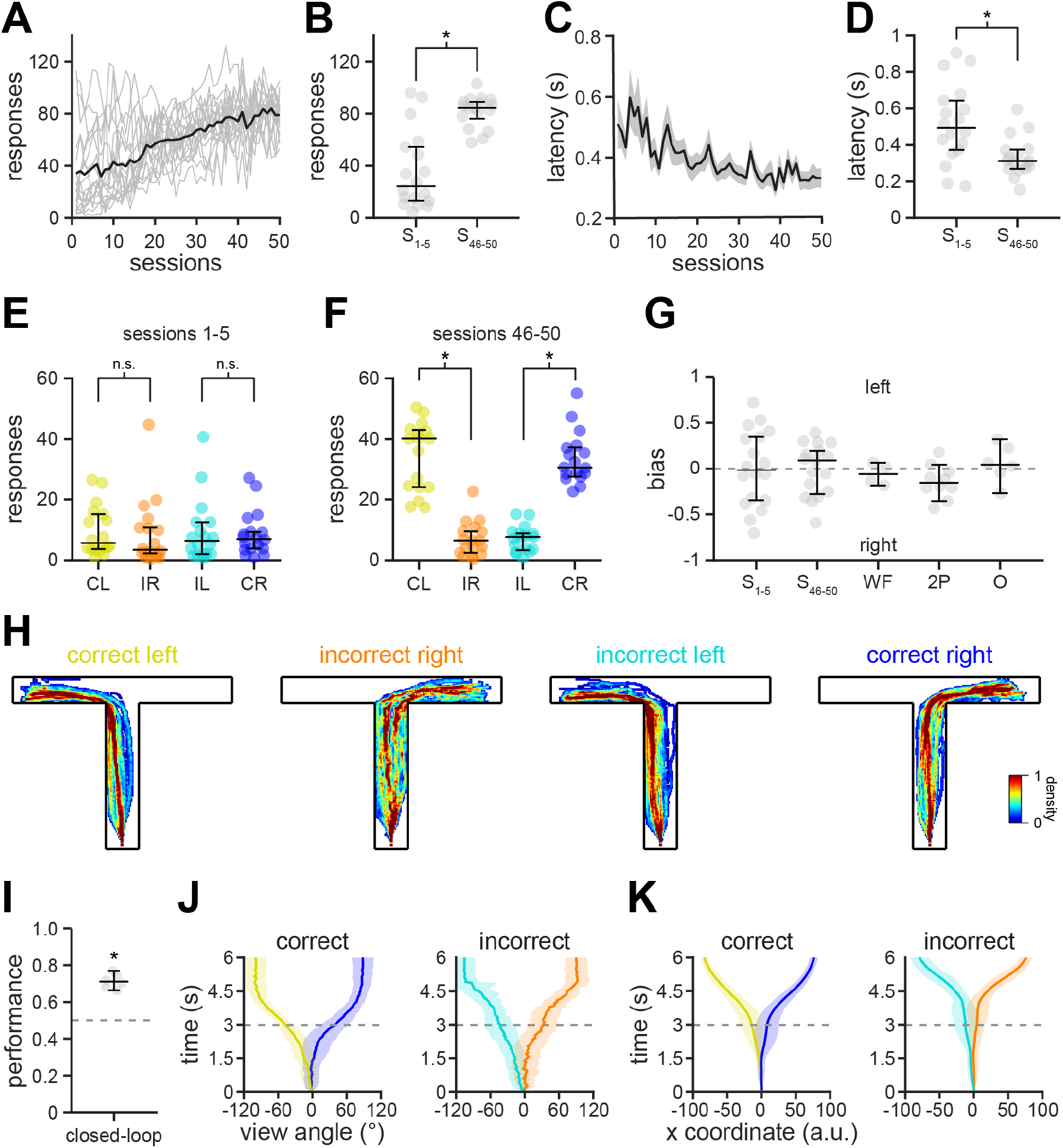
Behavioral performance for the context-trajectory association task. Related to Figure 1. **A**. Response count as a function of session during training (n = 18 mice). Grey lines: response count of individual mice; black line: average response count across mice. **B**. Scatter plots displaying response number of individual mice (grey dots) during the first (S_1-5_) and the last 5 days of training (S_46-50_). Mice significantly increased response number by the end of training (S_1-5_ = 24.2, 95% CI = [13, 54], S_46-50_ = 83.6, 95% CI = [73.1, 88.6]; *p* = 0.0002; Wilcoxon rank-sum test;). Black overlay: Median ± 95% CI. **C**. Response latency across training (*n* = 18 mice; mean ± s.e.m.). **D**. Scatter plots showing a decrease in response latency by the end of training (S_1-5_ = 489 ms, 95% CI = [370 ms, 638 ms], S_46-50_ = 309 ms, 95% CI = [267 ms, 371 ms]; *p* = 0.0048; Wilcoxon rank-sum test;). Individual mice are represented by grey dots. Black overlay: Median ± 95% CI. **E-F**. Fraction of correct and incorrect responses for both contexts at the beginning (CL_1-5_ = 5.6, 95% CI = [3.6, 15]; IR_1-5_ = 3.4, 95% CI = [2.2, 10.6],; *p* = 0.1891; IL_1-5_ = 6.3, 95% CI = [2, 12.4]; CR_1-5_ = 6.9, 95% CI = [3.8, 9.2]; *p* = 0.8123; Wilcoxon rank-sum test;) and at the end (CL_46-50_ = 39.8, 95% CI = [23.8, 42.2]; IR_46-50_ = 6.4, 95% CI = [2, 9.4]; *p* = 1.1×10^−6^; IL_46-50_ = 7.6, 95% CI = [3.4, 8.6]; CR_46-50_ = 30.4, 95% CI = [28.3, 36.8]; *p* = 7.0×10^−7^; Wilcoxon rank-sum test;) of training. Black overlay: Median ± 95% CI. **G**. Response bias calculated as the difference between all left responses and all right responses divided by the sum of all responses. Individual mice are represented by grey dots. No significant difference was found throughout training (S_1-5_ = −0.0177, 95% CI = [−0.3487, 0.3466]; S_46-50_ = 0.0897, 95% CI = [−0.2819, 0.1975]), widefield imaging (WF = −0.0635, 95% CI = [−0.1876, 0.0652]) or 2-photon imaging (2P = − 0.1580, 95% CI = [−0.2092, 0.0428]), and a slight left bias during optogenetics (O = 0.2598, 95% CI = [0.0613, 0.3256]) experiments. Data are considered significant if 95% confidence intervals are ≠ 0. Black overlay: Median ± 95% CI. **H**. Heat maps representing the position of mice (*n* = 45 sessions in 6 mice) in the virtual T-maze during closed-loop experiments, in which mice were able to turn the angle of view throughout the trial (see Methods). **I**. Behavioral performance during closed-loop experiments. Individual sessions are represented by grey dots (*n* = 45 sessions in 6 mice; closed-loop performance = 71.1, 95% CI = [66.3%, 76.9%], significant if lower 95% CI > 50%). Black overlay: Median ± 95% CI. **J-K**. View angle (J) and movement across the dimension parallel to the arms of the T-maze (K). Note that mice start turning towards the eventual choice prior to the decision point (mean ± s.d.).

**Figure S3.**
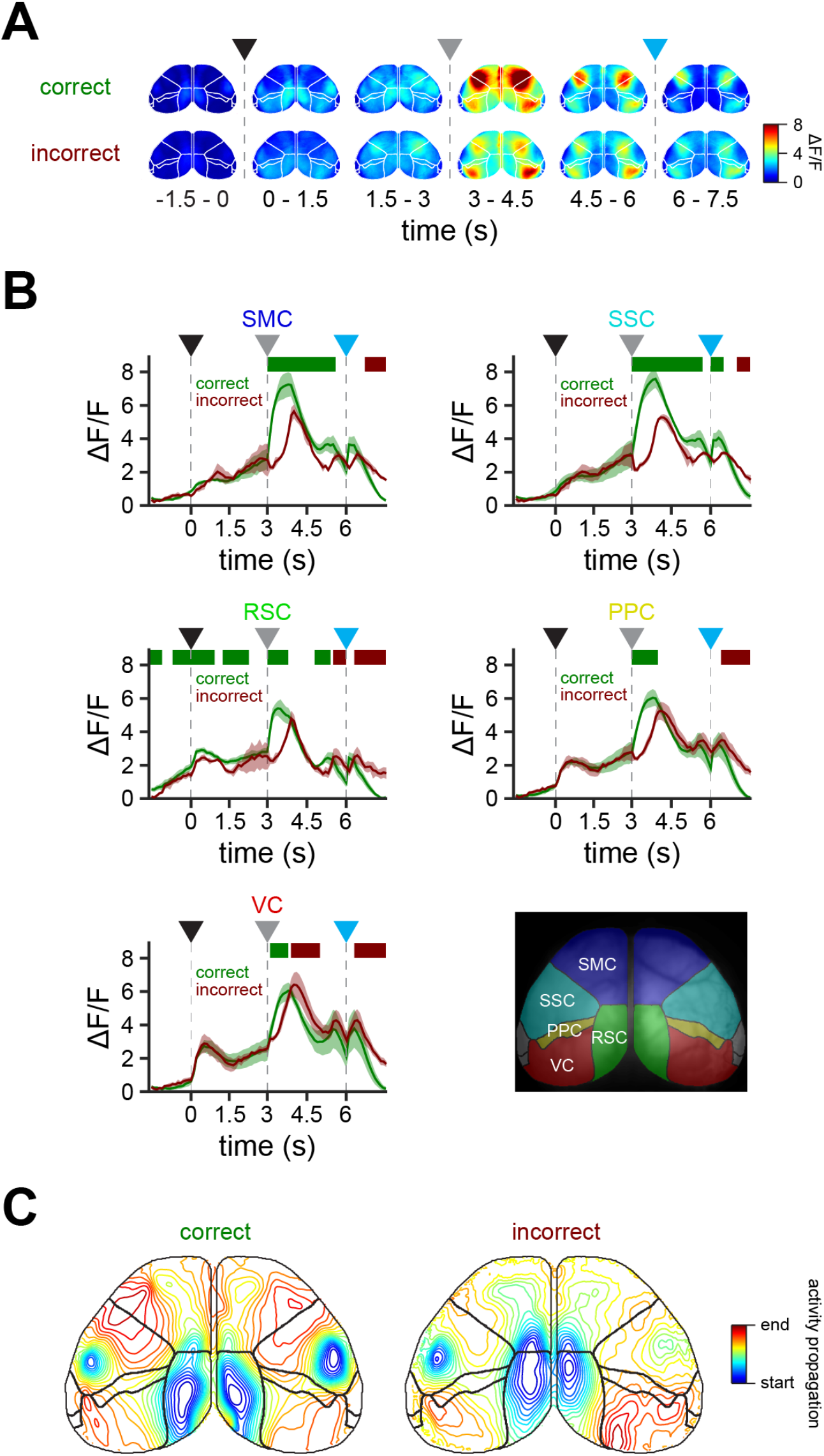
Mesoscale activity in dorsal cortex during context-trajectory associations. Related to Figure 2. **A**. Activity maps averaged across correct (top) and incorrect (bottom) trials (n = 20 sessions in 4 mice). **B**. Traces (mean ± s.e.m.) representing activity for correct (green) and incorrect (red) trials in somatomotor (SMC), somatosensory (SSC), retrosplenial (RSC), posterior parietal (PPC) and visual (VC) cortices (n = 20 sessions in 4 mice). Top green and red dashes indicate significantly higher activity in correct and incorrect trials, respectively (Wilcoxon rank-sum test). Significance only indicated for 5 or more consecutive samples. Bottom right, location of the regions of dorsal cortex imaged during performance of the task (SMC = somatomotor cortex, SSC = somatosensory cortex, RSC = retrosplenial cortex, PPC = posterior parietal cortex and VC = visual cortex). **C**. Phase maps (n = 20 sessions in 4 mice) showing the progression of activity in correct and incorrect trials during behavioral performance. Note that RSC is particularly important during the start of the trial, from where activity propagates towards premotor cortex (M2) and subsequently to caudal forelimb area (CFA). The propagation of activity exhibits an altered profile on incorrect trials. Arrowheads indicate trial start (black), decision point (grey) and trial end (cyan).

**Figure S4.**
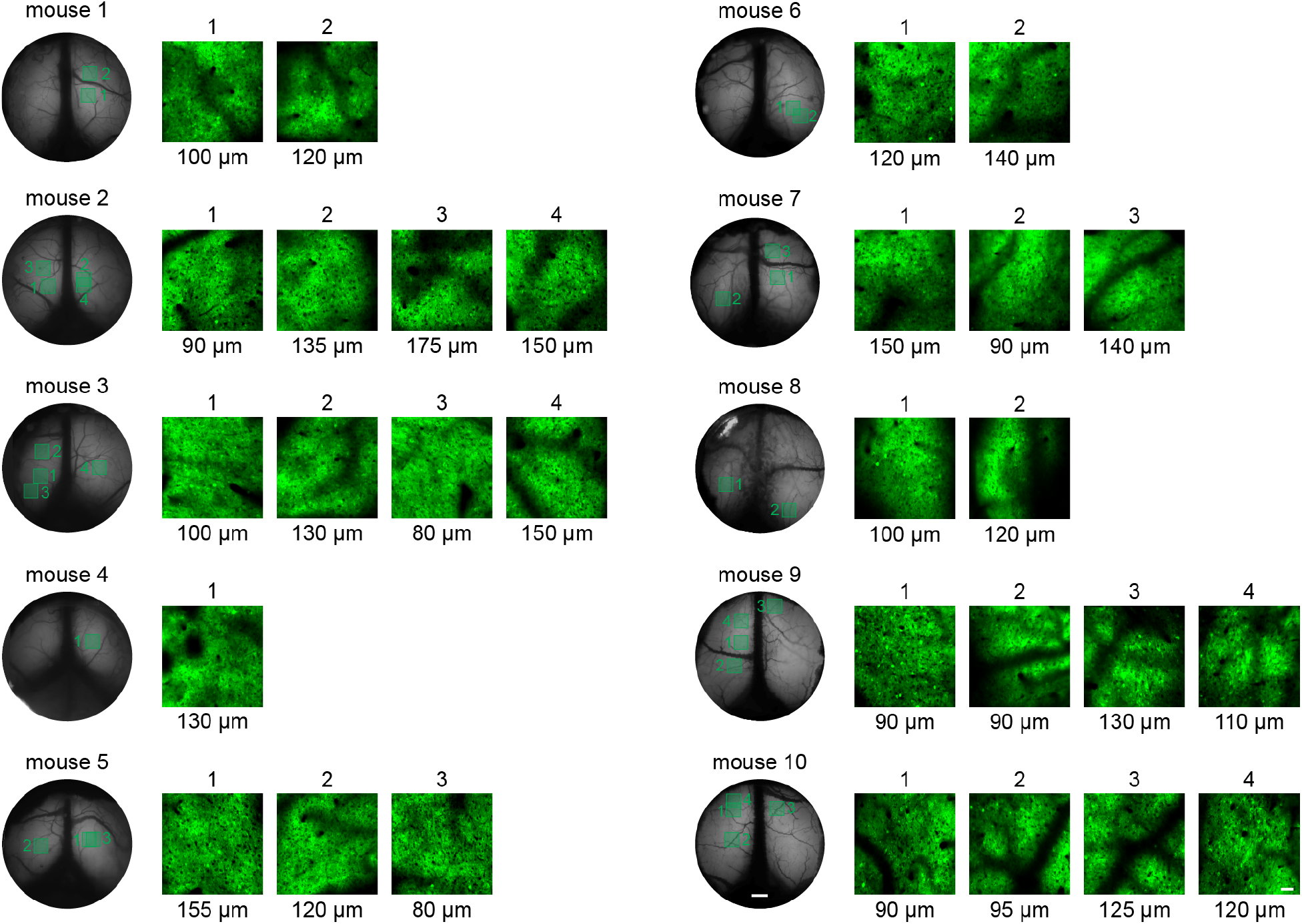
All 2-photon imaging fields and position in RSC. Related to Figure 4. Imaging fields included in 2-photon microscopy analysis: left, mean fluorescence of 4 mm cranial windows imaged with widefield microscopy (scale bar = 500 μm); right, mean fluorescence of 425 x 425 μm fields imaged with 2-photon microscopy (scale bar = 50 μm).

**Figure S5.**
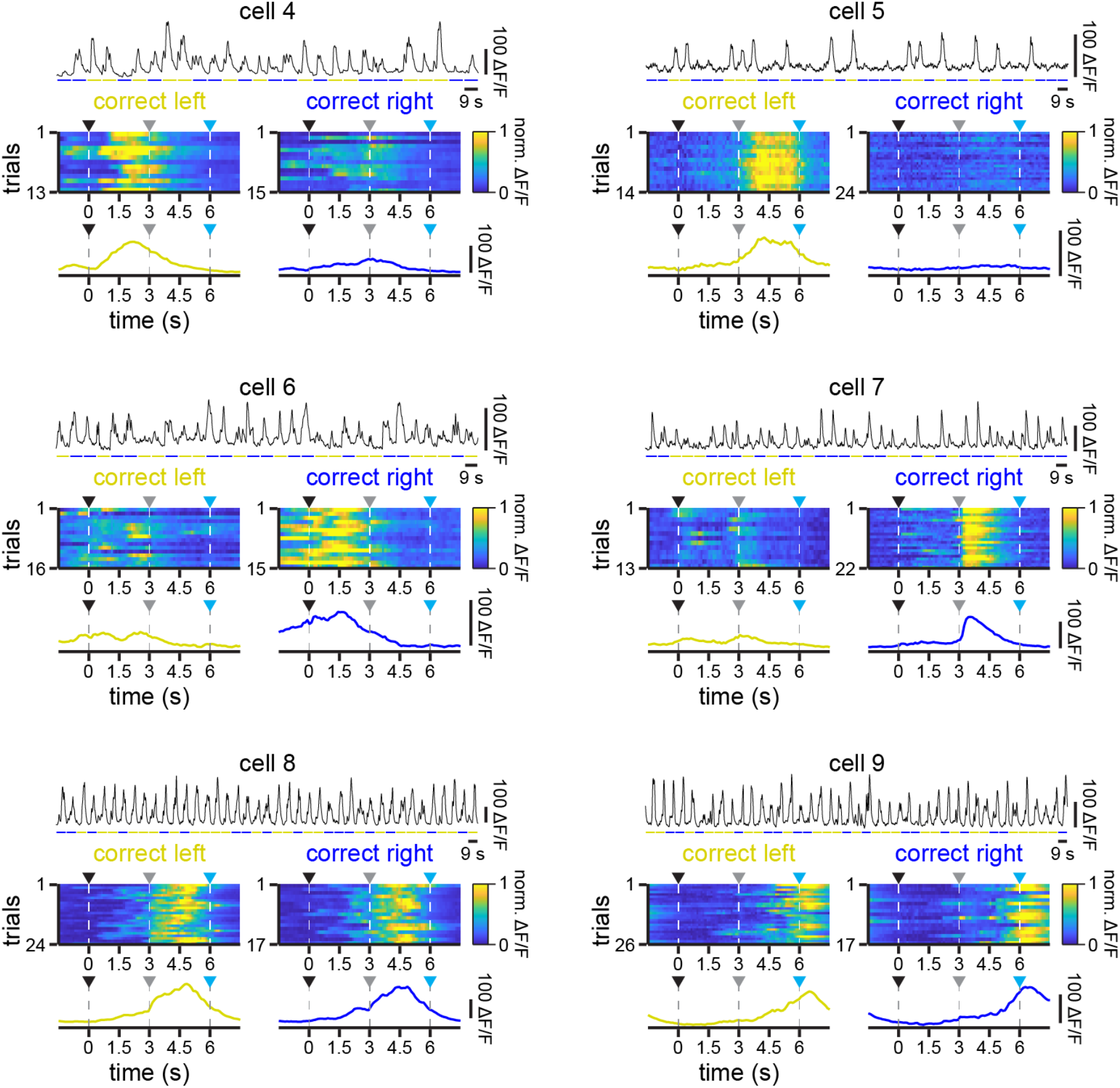
Example RSC neurons with significant responses during the task. Related to Figure 4. Additional example cells exhibiting significant responses to a particular context (cells 4 to 7) or to both contexts (cells 8 and 9). For each cell: Top, ΔF/F trace across concatenated correct trials for context 1 (yellow) and context 2 (blue). Middle, normalized responses to all correct trials for context 1 (left) and context 2 (right). Bottom, average activity across correct context 1 (left, yellow) and context 2 (right, blue) trials.

**Figure S6.**
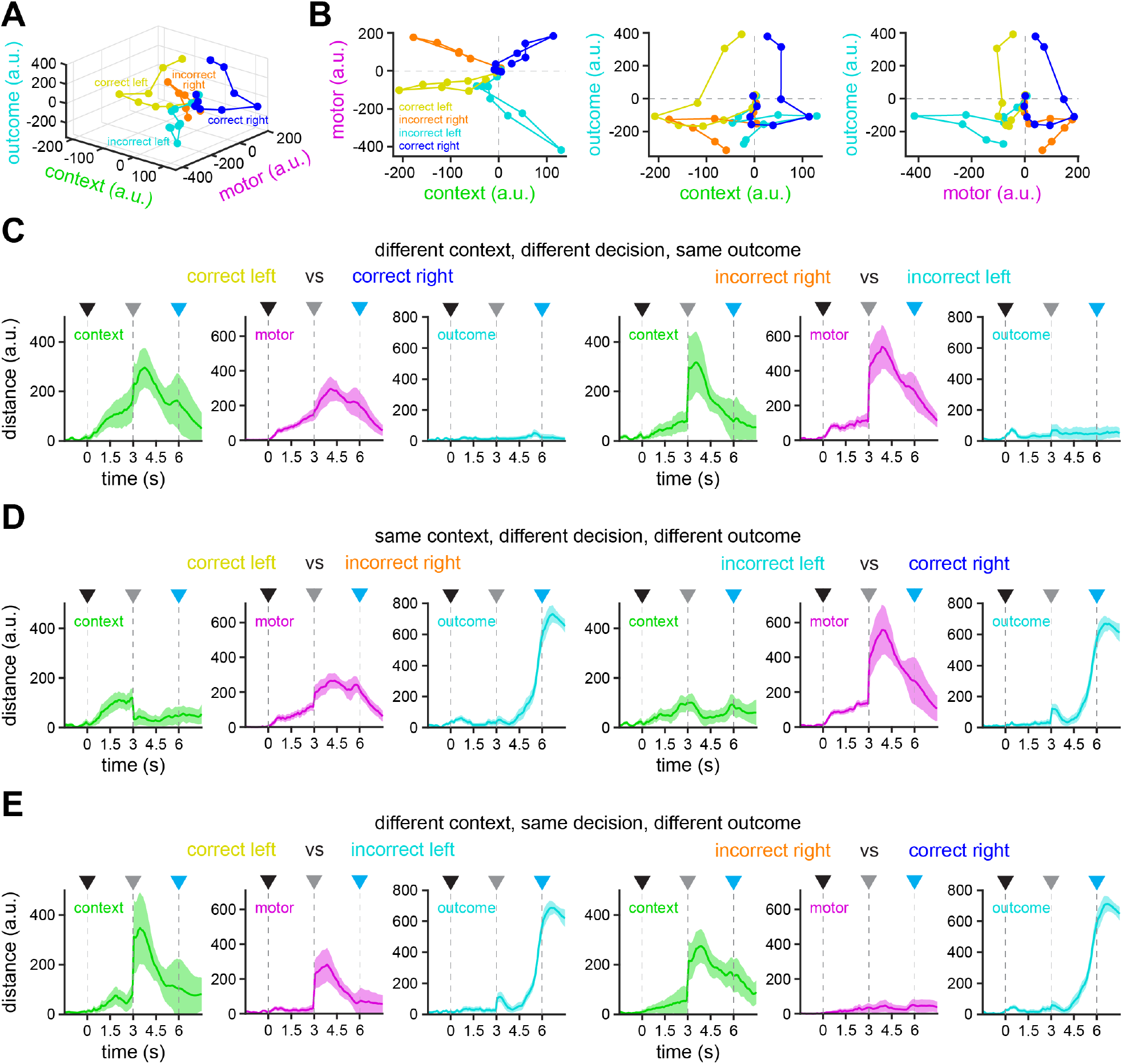
Population RSC activity in reduced dimensional space. Related to Figure 4. **A-B**. Trajectories of neuronal activity in correct and incorrect trials for both contexts obtained by targeted dimensional reduction for all task variables together (A) and for pairs of two variables (B). Note that activity is initially similar in each particular trial type, however, as trials progresses, activity diverges in relation to their context (context 1, negative; context 2, positive), motor decision (left, negative; right, positive) and outcome (no reward, negative; reward, positive). **C-E**. Distance between trajectories described by neuronal activity in the corresponding trials labeled context, motor and outcome (mean ± bootstrap-estimated s.e.m.). Arrowheads indicate trial start (black), decision point (grey) and trial end (cyan).

**Figure S7.**
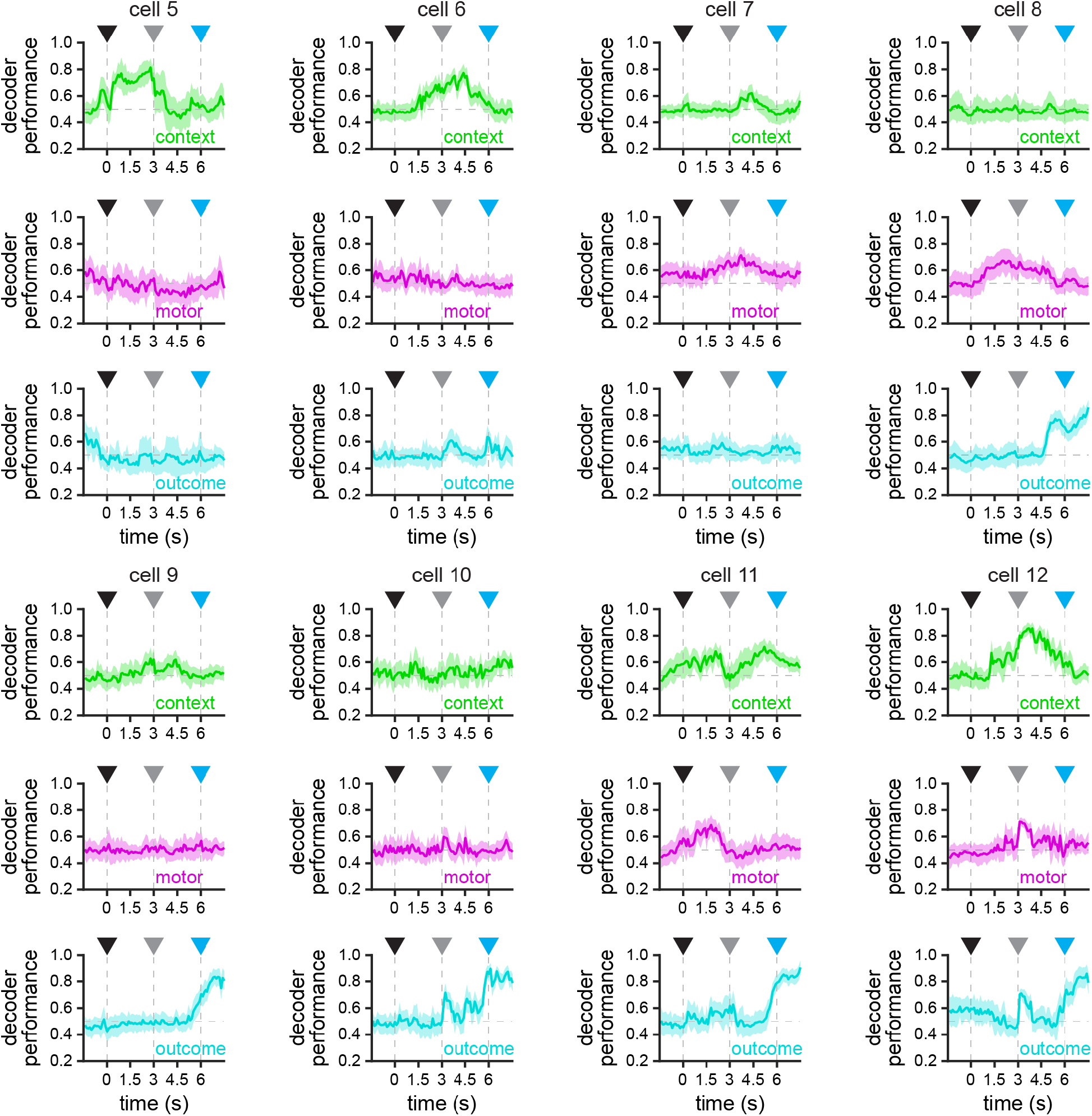
Encoding of task variables by single RSC neurons. Related to Figure 5. Additional example cells exhibiting significant decoding for context (cells 5 and 6), motor (cells 7 and 8), outcome (cells 9 and 10) or multiple (cell 11 and 12) task variables obtained by the support vector machine decoder (mean ± bootstrap-estimated s.e.m.).

**Figure S8.**
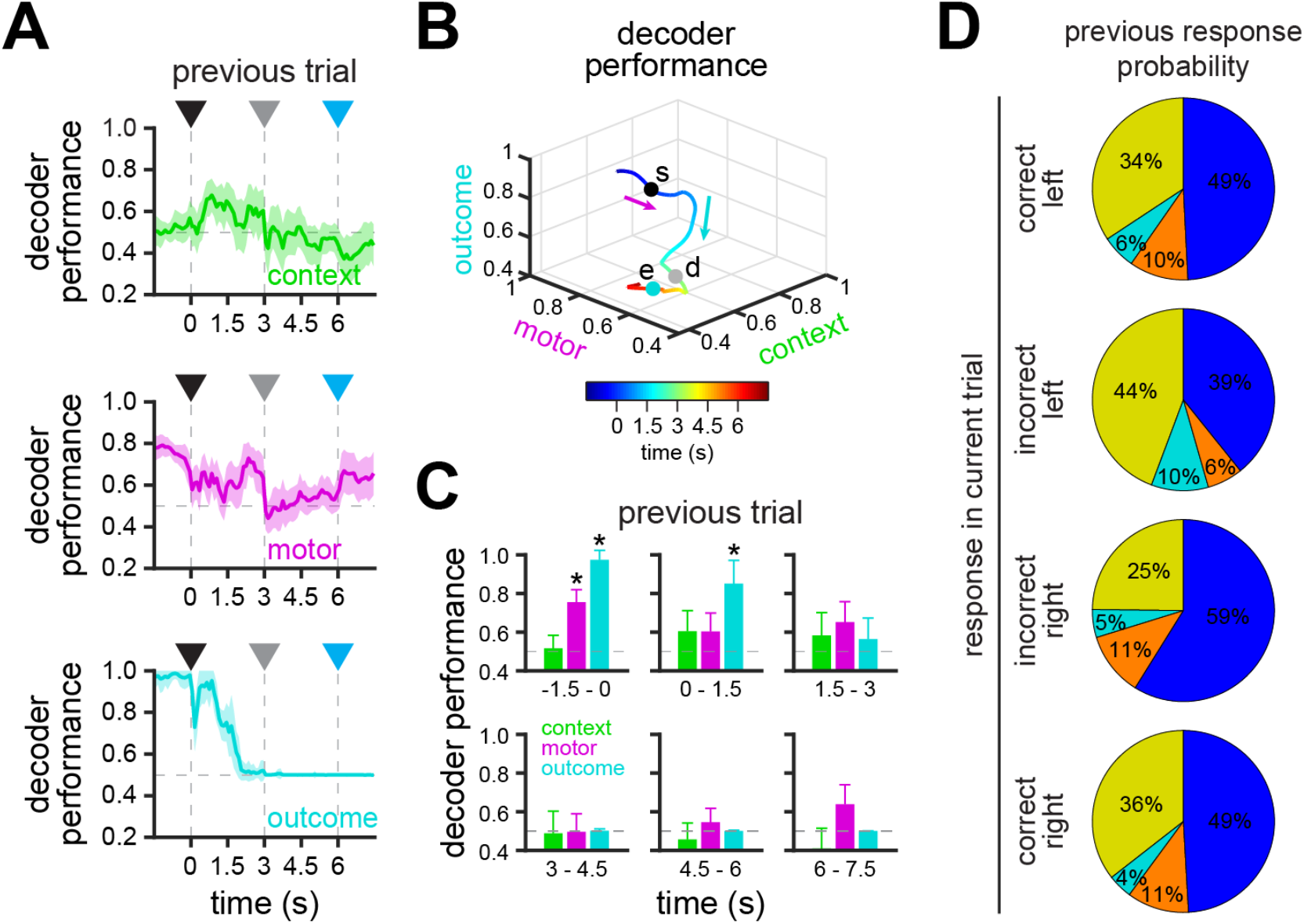
Encoding of task variables in the previous trial based on activity in the current trial. Related to Figure 5. **A**. Decoding of the context (top), motor (middle) or outcome (bottom) of previous trials based on the activity of current trials (n = 2,257 neurons) obtained by support vector machine (mean ± bootstrap-estimated s.e.m.). **B**. Trajectory described by population decoding of context, motor and outcome dimensions in previous trials (s, trial start; d, decision point; e, trial end). Note that decoding of motor and outcome in previous trials is high in the epoch corresponding to pre-trial activity, which decays following trial onset. Context decoding in previous trials is low throughout the trial. **C**. Bar plots showing significant decoding of previous trials of the different task variables in different epochs (n = 2,257 neurons; asterisks: lower 5% confidence interval > 0.5; mean ± bootstrap-estimated s.e.m.). **D**. Probability of correct and incorrect behavioral responses in the previous trial based on responses in the current trial (n = 29 sessions in 10 mice; yellow = correct left, cyan = incorrect left, orange = incorrect right, blue = correct right). Arrowheads indicate trial start (black), decision point (grey) and trial end (cyan).

**Figure S9.**
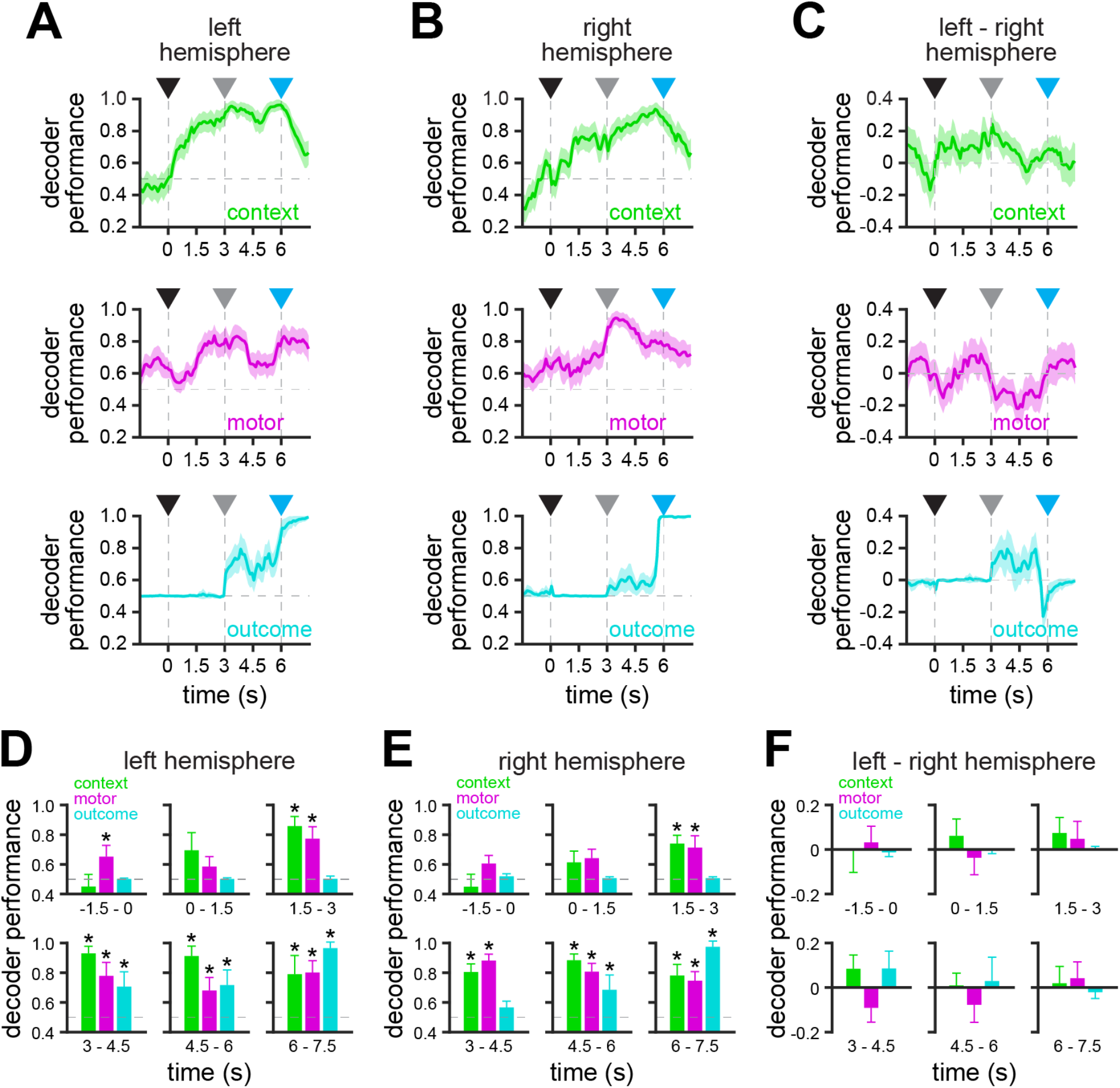
Encoding of task variables in each hemisphere. Related to Figure 6. **A-B**. Decoding of context (top), motor (middle) or outcome (bottom) in left (A) and right (B) hemispheres (left, *n* = 2,573 neurons; right, *n* = 2,621 neurons) obtained with population support vector machine decoder (mean ± bootstrap-estimated s.e.m.). **C**. Subtraction of performance of the different task variables in left hemisphere minus performance in right hemisphere (mean ± bootstrap-estimated s.e.m.). **D-E**. Bar plots showing significant decoding in left (D) and right (E) hemispheres of the different task variables in different epochs (left, *n* = 2,573 neurons; right, *n* = 2,621 neurons; asterisks: lower 5% confidence interval > 0.5; mean ± bootstrap-estimated s.e.m.). **F**. Bar plots showing the difference between decoding in left and right hemispheres of the different task variables in different epochs (mean ± bootstrap-estimated s.e.m.). None of the differences were significantly different. Arrowheads indicate trial start (black), decision point (grey) and trial end (cyan).

**Figure S10.**
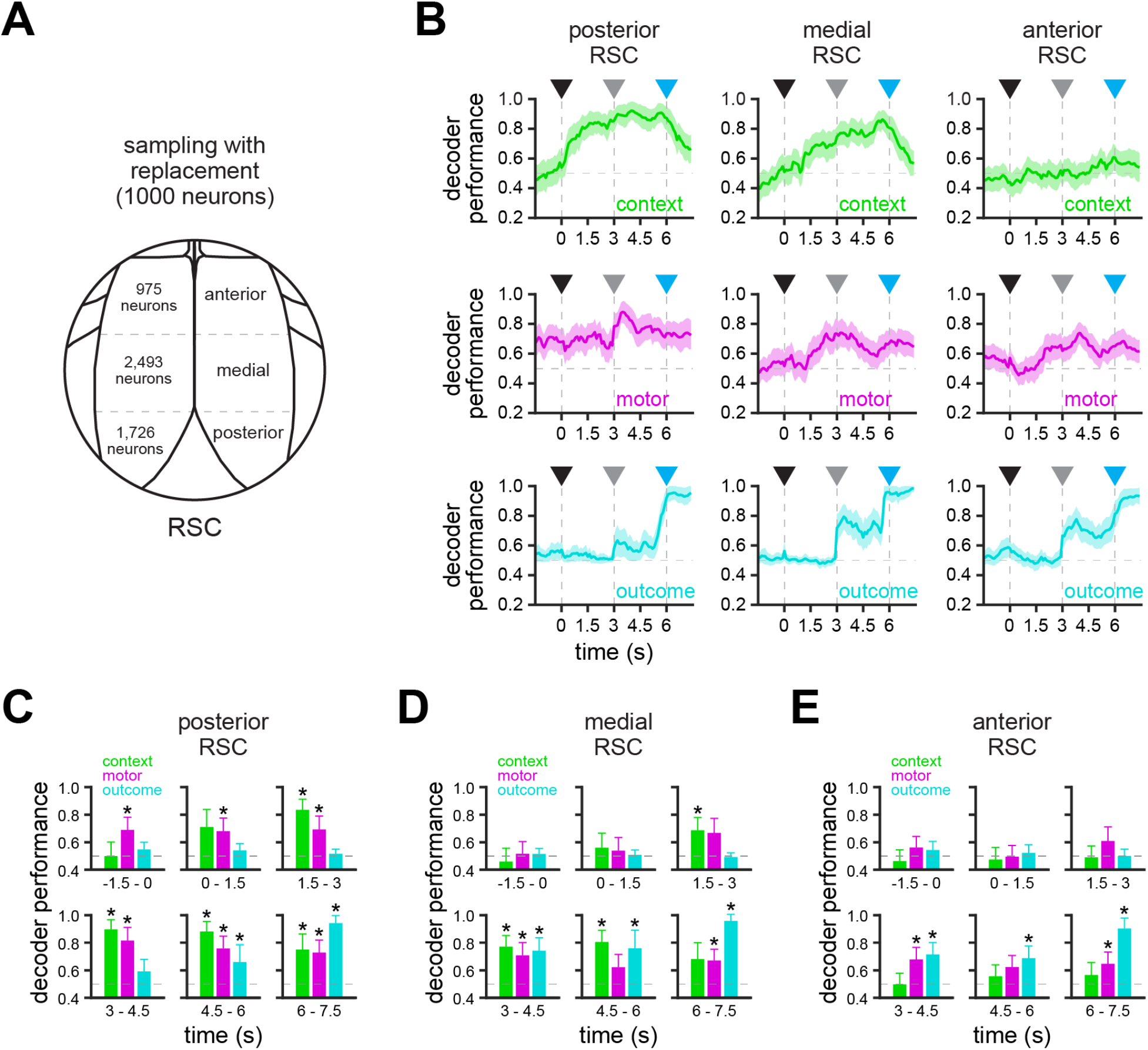
Encoding of task variables by anatomical location in RSC. Related to Figure 6. **A**. To control for the number of neurons recorded in different regions of RSC, encoding of the different task variables was calculated by sampling 1,000 neurons with replacement in each region. **B**. Decoding of context (top), motor (middle) and outcome (bottom) by populations of 1,000 neurons (sampled with replacement) in posterior, medial and anterior RSC (mean ± bootstrap-estimated s.e.m.). **C-E**. Bar plots showing significant decoding in posterior (C), medial (D) and anterior (E) RSC of context, motor and outcome in different epochs obtained by sampling 1,000 neurons with replacement (lower 5% confidence interval > 0.5; mean ± bootstrap-estimated s.e.m.). Note that using fixed population size does not impact results (see **Figure 6B-E**). Namely, context is preferentially encoded in posterior RSC, whereas motor and outcome decoding are more distributed across RSC but with different spatial and temporal dynamics. Arrowheads indicate trial start (black), decision point (grey) and trial end (cyan).

**Video S1. Example performance of context-to-trajectory association task**.

This clip shows five consecutive trials from a trained mouse performing the context-trajectory association task.

**Video S2. Encoding of task variables throughout RSC**.

Maps showing contribution of individual RSC neurons (*n* = 5,194) to the encoding of context (left), motor (middle), and outcome (right) task variables as a function of time throughout the trial. Some cells are saturated in order to display dynamic range.

## Notes

### Competing Interest Statement

The authors have declared no competing interest.

